# Angulin-1 seals tricellular contacts independently of tricellulin and claudins

**DOI:** 10.1101/2020.10.02.323378

**Authors:** Taichi Sugawara, Kyoko Furuse, Tetsuhisa Otani, Mikio Furuse

## Abstract

Tricellular tight junctions (tTJs) are specialized tight junctions (TJs) that seal the intercellular space at tricellular contacts (TCs), where the vertices of three epithelial cells meet. Tricellulin and angulin family membrane proteins are known constituents of tTJs, but the molecular mechanism of tTJ formation remains elusive. Here, we investigated the roles of angulin-1 and tricellulin in tTJ formation in MDCK II cells by genome editing. Angulin-1-deficient cells lost the plasma membrane contact at TCs with impaired epithelial barrier function. The COOH-terminus of angulin-1 bound to the TJ scaffold protein ZO-1 and disruption of their interaction influenced the localization of claudins at TCs, but not the tricellular sealing. Strikingly, the plasma membrane contact at TCs was formed in tricellulin- or claudin-deficient cells. These findings demonstrate that angulin-1 is responsible for the plasma membrane seal at TCs independently of tricellulin and claudins.

## Introduction

Epithelia work as barriers to separate the internal body from the external environment and to generate distinct fluid compartments within the body for various organ functions. Tight junctions (TJs) restrict leakage of solutes through the paracellular pathway, contributing to the epithelial barrier function (Tsukita et al., 2001)(Van Itallie and Anderson, 2014)(Zihni et al., 2016). On ultrathin section EM, TJs appear as a series of apparent fusions between adjacent cell membranes that obliterate the intercellular space at the most apical part of the lateral membrane (Farquhar and Palade, 1963). On freeze-fracture EM, TJs are visualized as anastomosing intramembranous particle strands (TJ strands) (Staehelin, 1973). Claudin family membrane proteins play key roles in TJ formation. Claudins assemble in cell-cell contacts between adjacent cells and polymerize within the plasma membrane to form TJ strands (Furuse et al., 1999)(Sasaki et al., 2003). On the cytoplasmic side of TJs, the claudin C-termini directly bind to three TJ-scaffolding proteins, ZO-1, ZO-2, and ZO-3 (Itoh et al., 1999). These three proteins have domain structures that include three PDZ domains (PDZ1–3) for protein-protein interactions (Willott et al., 1993)(Itoh et al., 1993). ZO-1 directly binds to claudins and junctional adhesion molecule-A (JAM-A) via its PDZ1 and PDZ3 domains, respectively (Itoh et al., 1999)(Itoh et al., 2001), and to ZO-2 via its PDZ2 domain (Fanning et al., 1998). Importantly, ZO-1 and ZO-2 are required for claudin-based TJ formation in cultured epithelial cells (Umeda et al., 2006)(Phua et al., 2014)(Otani et al., 2019).

As well as the paracellular barrier formed by TJs between adjacent cells, epithelial cells need to obliterate the extracellular space at tricellular contacts (TCs), where the vertices of three cells meet, to establish full epithelial barrier function. TCs contain specialized TJs designated tricellular TJs (tTJs) (Staehelin et al., 1969)(Staehelin, 1973)(Wade and Karnovsky, 1974)(Walker et al., 1985)(Ikenouchi et al., 2005). In freeze-fracture replicas, the most apical elements of TJ strands in bicellular TJs join together at TCs, turn, and extend in the basal direction attached to one another (Staehelin, 1973). Each vertical TJ strand along the apicobasal axis is called a central sealing element. Three central sealing elements at a TC squeeze the extracellular space to form a central tube of ~10-nm diameter, considered to work as a diffusion barrier (Staehelin, 1973). Short TJ strands are often connected to the central sealing elements in freeze-fracture replicas (Staehelin, 1973). Tricellulin and angulin family proteins, including angulin-1/LSR, angulin-2/ILDR1, and angulin-3/ILDR2, have been identified as molecular constituents of tTJs (Ikenouchi et al., 2005)(Masuda et al., 2011)(Higashi et al., 2013). Tricellulin is a four-transmembrane protein with structural similarity to occludin (Ikenouchi et al., 2005), while angulins are type I transmembrane proteins with a single Ig-like domain (Masuda et al., 2011)(Higashi et al., 2013). Tricellulin and angulins localize along the central sealing elements of tTJs (Ikenouchi et al., 2005)(Masuda et al., 2011). Because angulins recruit tricellulin to TCs (Masuda et al., 2011)(Higashi et al., 2013), the angulin-tricellulin axis is proposed to play crucial roles in tTJ formation (Furuse et al., 2014). RNAi-mediated suppression of tricellulin or angulin-1 expression in EpH4 mouse mammary epithelial cells was shown to impair epithelial barrier function (Ikenouchi et al., 2005)(Masuda et al., 2011). *In vivo* studies revealed that tricellulin or angulin-2 gene mutations cause human non-syndromic hearing loss (DFNB49 or DFNB42, respectively) (Riazuddin et al., 2006)(Borck et al., 2011), while tricellulin- or angulin-2-deficient mice show progressive hearing loss associated with hair cell degeneration (Kamitani et al., 2015)(Higashi et al., 2015)(Morozko et al., 2015)(Sang et al., 2015). Angulin-2-deficient mice also suffer from polyuria and polydipsia arising from renal concentrating defects (Gong et al., 2017). Meanwhile, angulin-1-deficient mouse embryos die before embryonic day 15.5 (Mesli et al., 2004) and exhibit blood-brain barrier leakage (Sohet et al., 2015).

Despite accumulating evidence on the physiological roles of tTJ-associated proteins, the molecular mechanism for tTJ formation remains to be solved. It is currently unclear whether the central sealing elements in tTJs contain TJ components like claudins, occludin, and ZO-1 because of the lack of detailed immunolocalization analyses. Furthermore, the molecules responsible for obliteration of the paracellular gap at TCs have not been clarified. To address these issues, it is useful to analyze the ultrastructure of tTJs in tricellulin- or angulin-deficient cultured epithelial cells like MDCK cells, which have contributed to investigations on structure-function relationships in TJs (Cereijido et al., 1978)(Stevenson et al., 1988)(Sonoda et al., 1999).

In this study, we examined the roles of angulin-1 and tricellulin in tTJ formation and epithelial barrier function by generating angulin-1- and tricellulin-deficient MDCK II cells using genome editing. We demonstrate that angulin-1 is required for the plasma membrane contact at TCs and recruits claudins to TCs through its interaction with ZO-1. We further show that the plasma membrane contact at TCs occurs independently of claudin-based TJ strands. Finally, we demonstrate that tricellulin is not essential for the plasma membrane contact at TCs or epithelial barrier function. Taken together, we conclude that angulin-1 plays pivotal roles in the plasma membrane seal at TCs for tTJ formation independently of tricellulin and claudins.

## Results

### Visualization of apicobasal extension of tTJs by fluorescence microscopy

To examine the detailed distributions of tTJ and TJ proteins at TCs, we performed immunofluorescence staining of frozen mouse kidney sections using Abs against tTJ and TJ proteins. Aquaporin-2 and claudin-2 were used as collecting duct and proximal tubule markers, respectively (Sabolić et al., 1995)(Kiuchi-Saishin et al., 2002). Upon close inspection of aquaporin-2-positive collecting duct epithelial cells, tricellulin was detected as rod-like signals extending from the intersection of TJ markers, including claudin-8, occludin, and ZO-1, i.e. TC, and appeared to merge with one of the branches of TJ markers extending in the basal direction (Fig. 1, A–C). Angulin-2 was colocalized with tricellulin (Fig. 1 D). Meanwhile, tricellulin was detected as dots at TCs in claudin-2-positive proximal tubule epithelial cells (Fig. 1 E). Next, we examined the distributions of tTJ and TJ proteins in MDCK II renal epithelial cells cultured on Transwell filters. In the image acquisition by confocal laser microscopy, we focused on confocal sections at two regions: apical region, in which TJ markers were highly concentrated, and lateral region, which was more basal than the apical region and did not contain TJ markers. The relationship between the structural organization of tTJs and the confocal sections is shown in Figure 2A. By immunofluorescence, tricellulin and angulin-1 were clearly detected at TCs in MDCK II cells, with relatively weak staining in the apical region and more intense staining in the lateral region (Fig. 2, B–E). In Z-stack confocal images, tricellulin and angulin-1 showed extended distributions along the apicobasal axis at TCs (Fig. 2 F). Furthermore, claudin-2, occludin, and ZO-1 were colocalized with tTJ markers along the apicobasal axis at TCs (Fig. 2, B–E). Taken together, these observations indicate that TJ proteins are incorporated into tTJs, the outlines of which can be visualized by light microscopy using tTJ markers.

**Figure 1.**
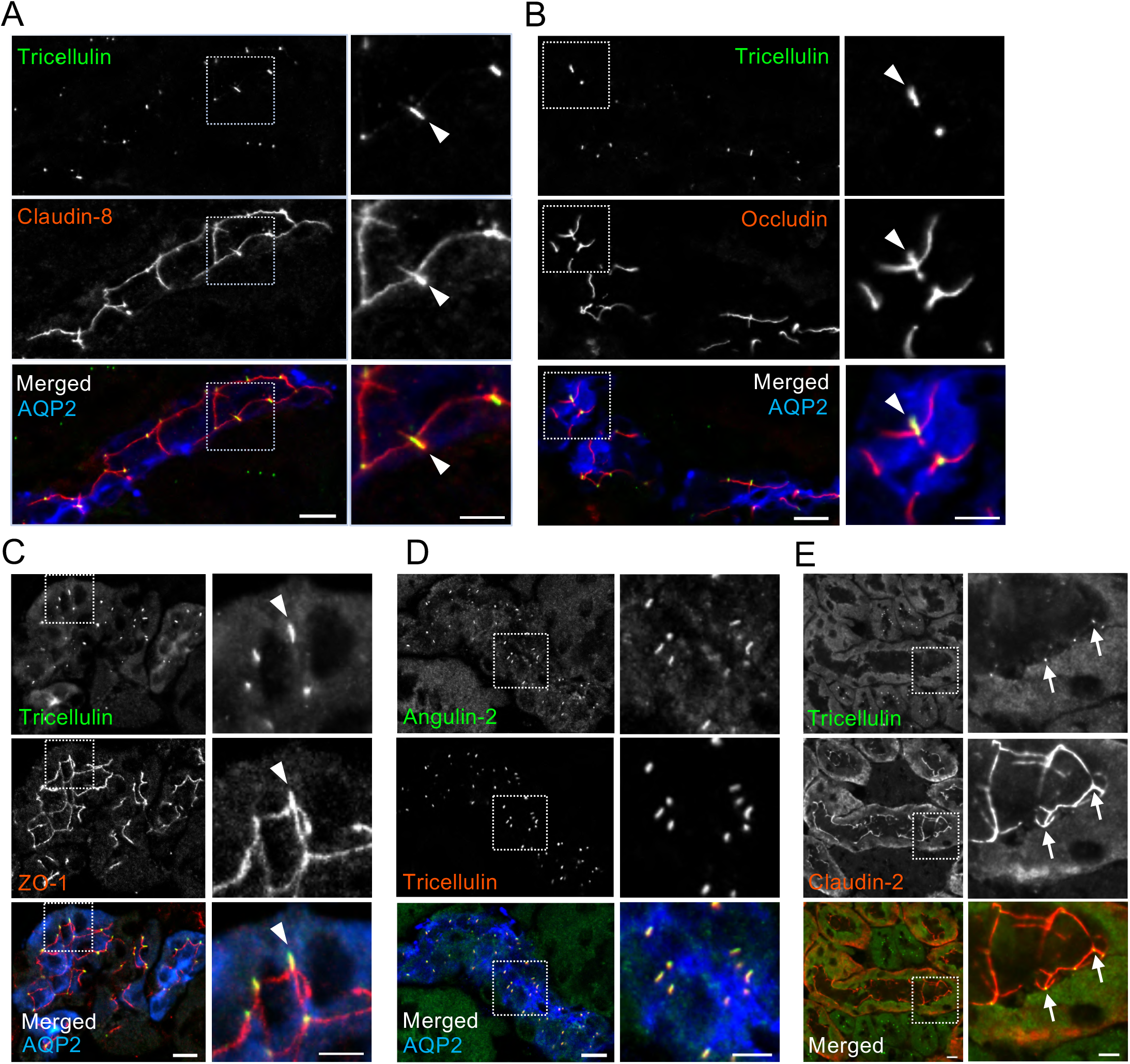
Localization of tTJ and TJ proteins in collecting ducts and proximal tubules in the mouse kidney. **(A–D)** Triple immunofluorescence staining of frozen mouse kidney sections containing collecting ducts with anti-tricellulin mAb, anti-claudin-8 pAb, and anti-AQP2 pAb (A), anti-tricellulin mAb, anti-occludin pAb, and anti-AQP2 pAb (B), anti-tricellulin mAb, anti-ZO-1 pAb, and anti-AQP2 pAb (C), and anti-angulin-2 pAb, anti-tricellulin mAb, and anti-AQP2 pAb (D). AQP2 staining is only shown in the merged images. The boxed regions are magnified on the right. Tricellulin shows rod-like staining in AQP2-positive collecting ducts. Claudin-8 (A), occludin (B), and ZO-1 (C) colocalize with tricellulin at TCs (arrowheads). **(E)** Double immunofluorescence staining of a frozen mouse kidney section containing proximal tubules with anti-tricellulin mAb and anti-claudin-2 pAb. The boxed regions are magnified on the right. In claudin-2-positive proximal tubules, tricellulin signals appear as dots at TCs (arrows). Bars: 10 μm (left), 5 μm (right).

**Figure 2.**
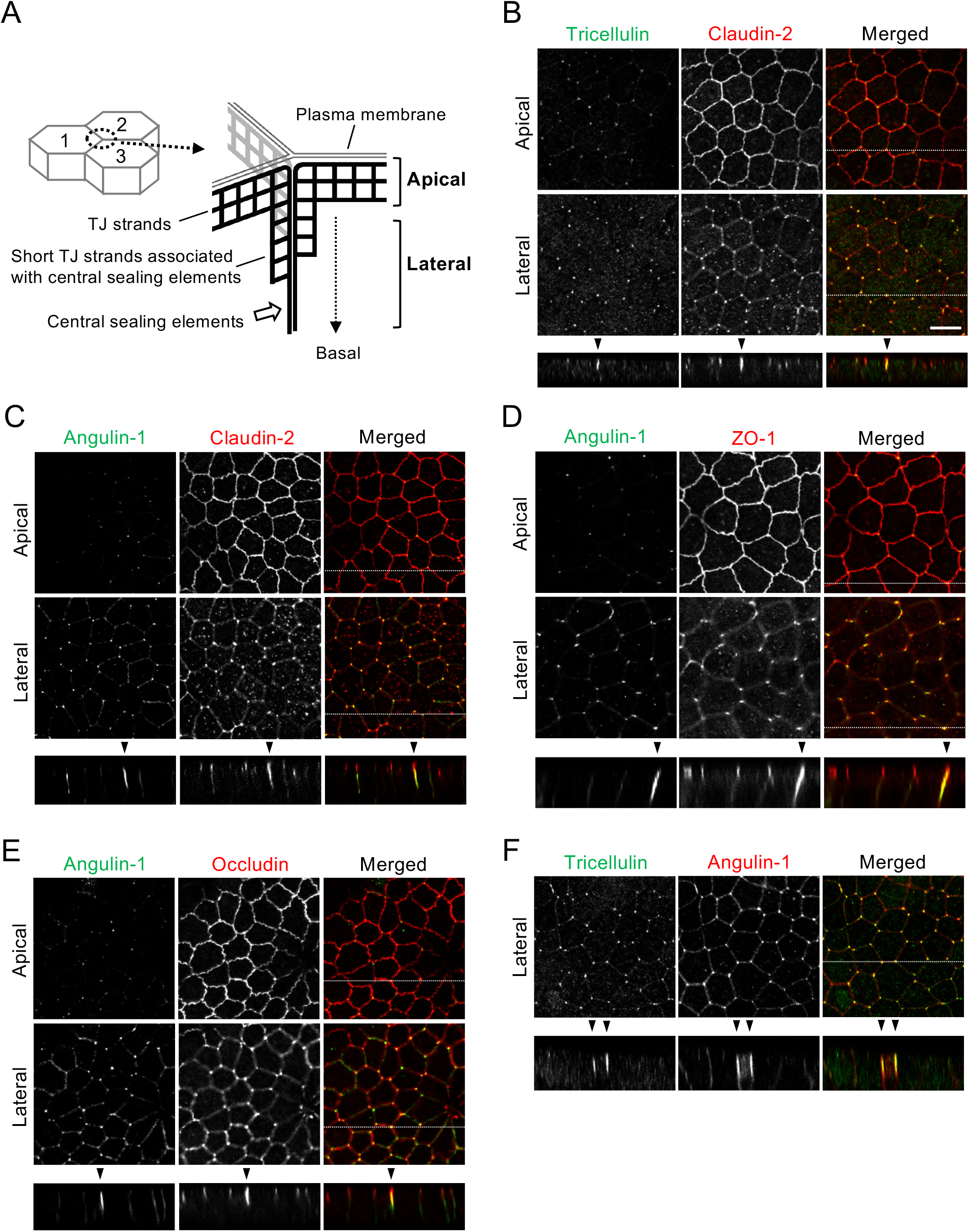
Localization of tTJ and TJ proteins at TCs in MDCK II cells. **(A)** Supposed relationship between tTJ structural organization and confocal sections in the apical and lateral regions. The scheme represents tTJs observed from the inside of Cell #3. The network of black bold lines represents TJ strands. The most apical elements of TJ strands in bicellular TJs join together at TCs, turn, and extend in the basal direction attached to one another to form the central sealing elements that connect with short TJ strands. The confocal sections in the apical region correspond to the level of bicellular TJs, while those in the lateral region are more basal and do not contain bicellular TJs. **(B–F)** Double immunofluorescence staining of MDCK II cells with anti-tricellulin pAb and anti-claudin-2 mAb (B), anti-angulin-1 pAb and anti-claudin-2 mAb (C), anti-angulin-1 pAb and anti-ZO-1 mAb (D), anti-angulin-1 pAb and anti-occludin mAb (E), and anti-tricellulin mAb and anti-angulin-1 pAb (F). Confocal sections in the apical region including TJ markers and the lateral region together with the corresponding Z-stack images along the white dotted lines are shown. Not only tricellulin and angulin-1, but also claudin-2, occludin, and ZO-1 show extended localization along the apicobasal axis at TCs (arrowheads). Bar: 10 μm.

### Angulin-1 is essential for the plasma membrane contact at TCs and epithelial barrier function in MDCK II cells

To examine the role of angulins in tTJ formation, we generated angulin-deficient MDCK II cells. Expression of angulin-1 and angulin-2 was previously demonstrated in MDCK cells by RNA sequencing analysis, with much higher expression of angulin-1 than angulin-2 (Shukla et al., 2015). Thus, we focused on angulin-1 in the present study. We disrupted the angulin-1-encoding gene in MDCK II cells by TALEN-mediated genome editing and established two independent angulin-1 knockout (KO) cell clones: angulin-1 KO_1 and angulin-1 KO_2 (Figs. 3 A and S1 A). Because the two clones had essentially the same phenotype, we present data for angulin-1 KO_1 cells as representatives of angulin-1 KO cells unless otherwise specified. Immunofluorescence staining revealed that TC localization of tricellulin was impaired in angulin-1 KO cells (Fig. 3 B), supporting the idea that angulin-1 is the major angulin subtype in MDCK II cells. In angulin-1 KO cells, claudin-2 and ZO-1 were detected in the apical region containing TJs, but not in the lateral region at TCs (Fig. 3 C). Re-expression of angulin-1 in angulin-1 KO cells restored the extended distributions of claudin-2 and ZO-1 along the apicobasal axis at TCs in Z-stack confocal images (Figs. 3 C and S1 B). These results suggest that angulin-1 is required for vertically extended localization of TJ proteins at TCs.

**Figure 3.**
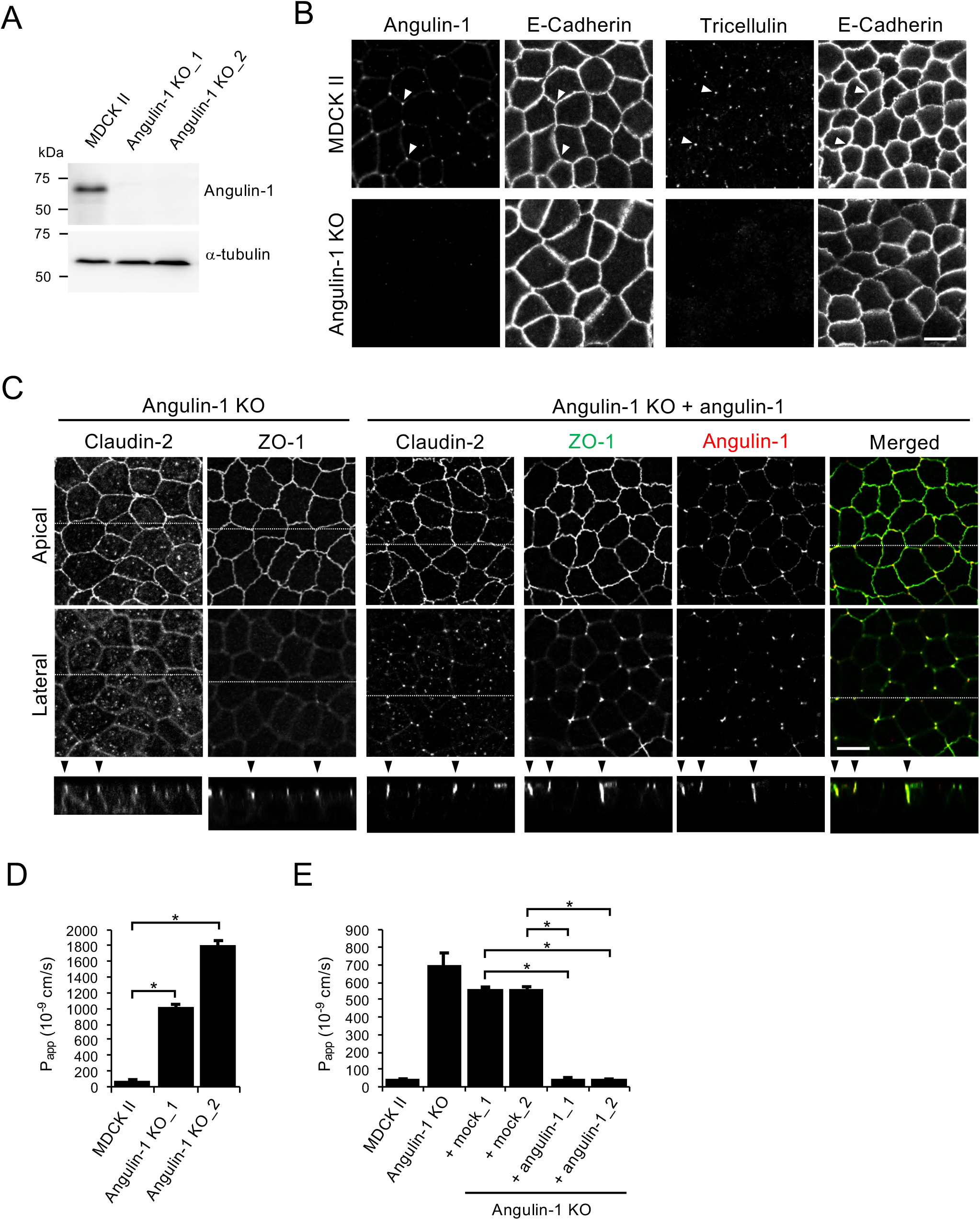
Role of angulin-1 in the localization of TJ proteins at TCs and epithelial barrier function. **(A)** Western blotting of lysates of MDCK II cells and two independent angulin-1 KO cell clones (angulin-1 KO_1 and KO_2) with anti-angulin-1 pAb or anti-α-tubulin mAb. **(B)** Double immunofluorescence staining of MDCK II cells and angulin-1 KO cells with anti-angulin-1 pAb and anti-E-cadherin mAb or anti-tricellulin pAb and anti-E-cadherin mAb. Arrowheads indicate TCs. **(C)** Immunofluorescence staining of angulin-1 KO cells and angulin-1 KO cells expressing exogenous angulin-1 (angulin-1 KO + angulin-1). Angulin-1 KO cells were immunostained with anti-claudin-2 mAb or anti-ZO-1 mAb. Angulin-1 KO cells expressing exogenous angulin-1 were singly immunostained with anti-claudin-2 mAb or doubly stained with anti-ZO-1 mAb and anti-angulin-1 pAb. Confocal sections of the apical region including TJ markers and the lateral region together with the corresponding Z-stack images along the white dotted lines are shown. Arrowheads indicate TCs. **(D)** Paracellular flux of fluorescein in MDCK II cells and two independent angulin-1 KO cell clones (angulin-1 KO_1 and KO_2). **(E)** Paracellular flux of fluorescein in MDCK II cells, angulin-1 KO cells, two independent mock-transfected angulin-1 KO cell clones (+mock_1 and +mock_2), and two independent angulin-1 KO cell clones expressing exogenous angulin-1 (+angulin-1_1 and +angulin-1_2). Data are shown as mean ± SD (*n*=3) and were analyzed by Dunnett’s test (D) or the Tukey–Kramer test (E). \**P*<0.01. Bars: 10 μm.

Next, we examined the epithelial barrier function in angulin-1 KO cells cultured on Transwell filters. Paracellular flux of fluorescein (332 Da) was increased in angulin-1 KO cells compared with MDCK II cells (Fig. 3 D), and the increase was abolished by re-expression of mouse angulin-1 (Fig. 3 E). Meanwhile, transepithelial electrical resistance (TER), reflecting electrolyte permeability, was not significantly altered in angulin-1 KO cells (Fig. S1 C). The low TER in MDCK II cells arising from endogenous claudin-2, which forms cation-selective pores in TJs (Furuse et al., 2001) (Amasheh et al., 2002)(Tokuda and Furuse, 2015), may hamper detection of subtle differences in TER measurements. This prompted us to further examine the impact of angulin-1 on electrolyte permeability. We disrupted the angulin-1 gene by CRISPR/Cas9-mediated genome editing in claudin-2-deficient MDCK II cells (claudin-2 KO cells) with high TER (Tokuda and Furuse, 2015) and established three angulin-1/claudin-2 double KO cell clones (Fig. S2, A and B). In all three clones, TER was remarkably decreased and paracellular flux of fluorescein was increased compared with control claudin-2 KO cells (Fig. S2 C and D). Re-expression of angulin-1 in angulin-1/claudin-2 double KO cells restored the TER and flux of fluorescein to the levels in claudin-2 KO cells (Fig. S2, E–G). These findings suggest that angulin-1 plays crucial roles in epithelial barrier formation in MDCK II cells.

To investigate the role of angulin-1 in tTJ ultrastructure, we prepared horizontal ultrathin sections of cells cultured on Transwell filters for transmission EM. Intriguingly, the extracellular space at TCs was obliterated by the contact between the three plasma membranes in MDCK II cells. This occurred at not only the level containing TJs (TJ level), but also a more basal level where desmosomes were observed (desmosome level) (Fig. 4 A). The structures appeared to correspond to the tTJs previously described by freeze-fracture EM (Staehelin, 1973). In contrast, no plasma membrane contact was observed at TCs in angulin-1 KO cells. Instead, a gap was observed at TCs in both the TJ and desmosome levels (Fig. 4 A). Re-expression of angulin-1 in angulin-1 KO cells restored the plasma membrane contact at TCs (Fig. 4 A). We further examined the tTJ ultrastructure by freeze-fracture EM. In MDCK II cells, typical tTJs with the central sealing elements and short TJ strands connected to these elements were observed (Fig. 4, B_1_, B_1_’, and B_2_). The depth of the tTJs varied. In contrast, TCs in angulin-1 KO cells had two vertical TJ strands separated from one another by smooth fracture planes of the plasma membranes (Fig. 4, B_3_, B_3_’, and B_4_). TJ strands along bicellular contacts were continuous in angulin-1 KO cells (Fig. 4 B_5_). These results demonstrate that angulin-1 is required for the plasma membrane contact at TCs and the central sealing elements of tTJs.

**Figure 4.**
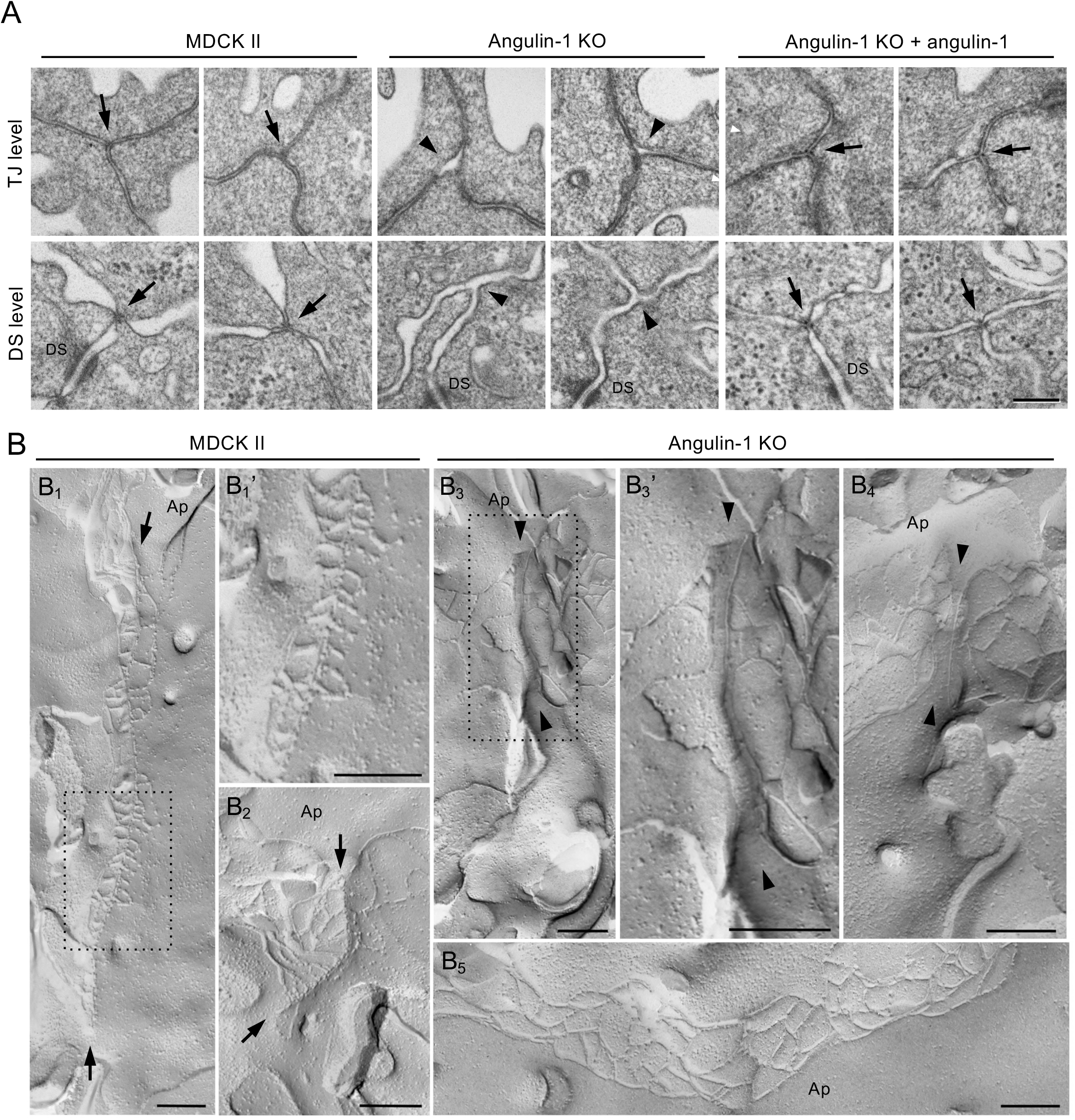
Angulin-1 is required for tTJ formation. **(A)** EM observation of horizontal ultrathin sections of MDCK II cells, angulin-1 KO cells, and angulin-1 KO cells expressing exogenous angulin-1 (angulin-1 KO + angulin-1). The three plasma membranes seal TCs at not only the level of TJs (TJ level), but also the more basal level of the lateral membrane containing desmosomes (DS level) in MDCK II cells and angulin-1 KO cells expressing exogenous angulin-1 (arrows). In contrast, a gap is observed at TCs in angulin-1 KO cells (arrowheads). DS: desmosome. **(B)** Freeze-fracture EM observation of TC regions in MDCK II cells and angulin-1 KO cells. In MDCK II cells (B_1_, B_1_’, and B_2_), attached central sealing elements formed by vertical TJ strands are observed (between arrows) with variation in length depending on the TCs. B_1_’ is a magnified image of the boxed region in B_1_. At TCs in angulin-1 KO cells (B_3_, B_3_’, and B_4_), attached central sealing elements are not found, but vertical TJ strands separated by flat planes of the plasma membranes are observed (between arrowheads). B_3_’ is a magnified image of the boxed region in B_3_. Continuous TJ strands are observed at bicellular contacts in angulin-1 KO cells (B_5_). Ap: apical side. Bars: 200 nm.

### PDZ domain-binding motif of angulin-1 is required for extended localization of TJ constituents at TCs

Angulin family proteins have a putative PDZ domain-binding motif (pbm) at their C-terminus (Higashi et al., 2013). However, it is unknown which molecules the pbm interacts with and how the putative interactions contribute to tTJ formation. To examine the role of the angulin-1 pbm, we established angulin-1 KO cells stably expressing an angulin-1 mutant lacking the C-terminal 5 amino acids (angulin-1 Δpbm) (Figs. 5 A and S1 B). On immunofluorescence staining, angulin-1Δpbm was localized at TCs and extended along the apicobasal axis with tricellulin (Fig. 5 B). Meanwhile, ZO-1 and claudin-2 were localized at the TJ level only, and did not colocalize with angulin-1 Δpbm along the apicobasal axis at TCs (Fig. 5 B). These results suggest that the angulin-1 pbm is responsible for the extended distribution of TJ constituents at TCs.

**Figure 5.**
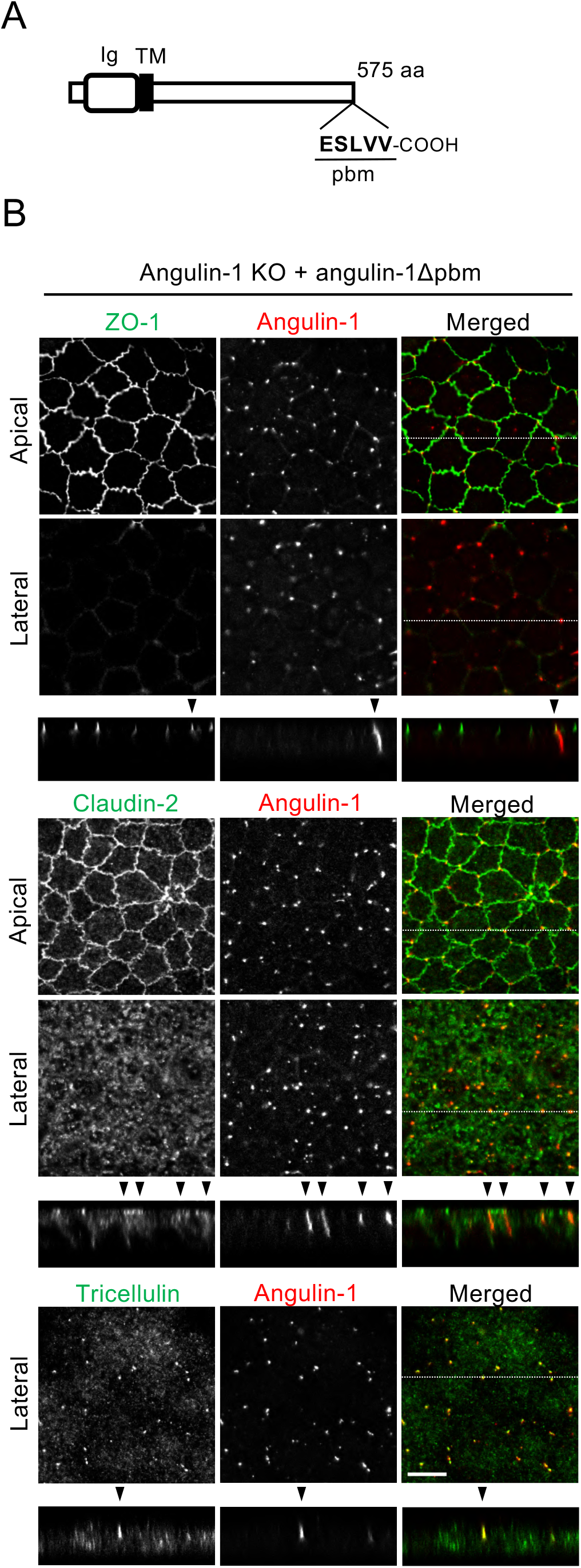
Role of the angulin-1 pbm in the localization of TJ proteins at TCs. **(A)** Schematic drawing of mouse angulin-1 with 575 amino acids. Five amino acids of the putative pbm at the C-terminus are shown. Ig: immunoglobulin-like domain; TM: transmembrane domain. **(B)** Double immunofluorescence staining of angulin-1 KO cells expressing exogenous angulin-1 lacking the C-terminal five amino acids (angulin-1 Δpbm) with anti-ZO-1 mAb and anti-angulin-1 mAb, anti-claudin-2 mAb and anti-angulin-1 mAb, or anti-tricellulin pAb and anti-angulin-1 mAb. Confocal sections of the apical region including TJ markers and the lateral region together with the corresponding Z-stack images along white dotted lines are shown. Arrowheads indicate TCs. Bar: 10 μm.

These observations prompted us to examine whether the angulin-1 pbm binds to the PDZ domains of ZO proteins. We generated bacterial GST-fusion proteins of the C-terminal 167 amino acids (aa 409–575) of mouse angulin-1 and its deletion mutant lacking the C-terminal 5 amino acids corresponding to the pbm (aa 409–570), designated GST-ang575 and GST-ang570, respectively. Pull-down assays revealed that GST-ang575, but not GST-ang570, interacted with ZO-1, ZO-2, and ZO-3 in angulin-1 KO cell lysates (Fig. 6 A), suggesting that the angulin-1 pbm binds to ZO family proteins. We also attempted co-immunoprecipitation experiments from MDCK II cells but were unsuccessful, possibly due to the poor solubility of TJ components (data not shown). Thus, we focused on ZO-1 as a possible binding partner of angulin-1. We performed pull-down assays with GST-ang575 and a bacterial maltose-binding protein (MBP)-fusion protein of aa 1–862 of mouse ZO-1 containing three PDZ domains (MBP-N-ZO-1) (Fig. 6 B). We found that GST-ang575, but not GST, bound to MBP-N-ZO-1 (Fig. 6 C). To determine which PDZ domain of ZO-1 binds to angulin-1, we generated bacterial MBP-fusion proteins of aa 19–113, aa 181–292, and aa 423–503 of mouse ZO-1, corresponding to PDZ1, PDZ2, and PDZ3, respectively (Fig. 6B). In pull-down assays, GST-ang575 bound to the MBP-fusion protein of PDZ2, but not to those of PDZ1 or PDZ3 (Fig. 6 D), suggesting that angulin-1 directly binds to PDZ2 of ZO-1 via its pbm. To determine the role of the binding between angulin-1 and ZO-1 in the localization of TJ proteins along the apicobasal axis at TCs, we expressed a chimeric protein of angulin-1 Δpbm and full-length ZO-1 (angulin-1 Δpbm-ZO-1) in angulin-1 KO cells (Fig. 6, E and F). Immunofluorescence staining of stable transfectants with an anti-angulin-1 pAb showed vertically extended localization of angulin-1 Δpbm-ZO-1. Moreover, claudin-2 was colocalized with angulin-1 Δpbm-ZO-1 along the apicobasal axis at TCs (Fig. 6 G), different from the case with angulin-1 Δpbm (Fig. 5 B). These results suggest that the interaction between angulin-1 and ZO-1 supports the extended localization of claudin-2 along the apicobasal axis at TCs.

**Figure 6.**
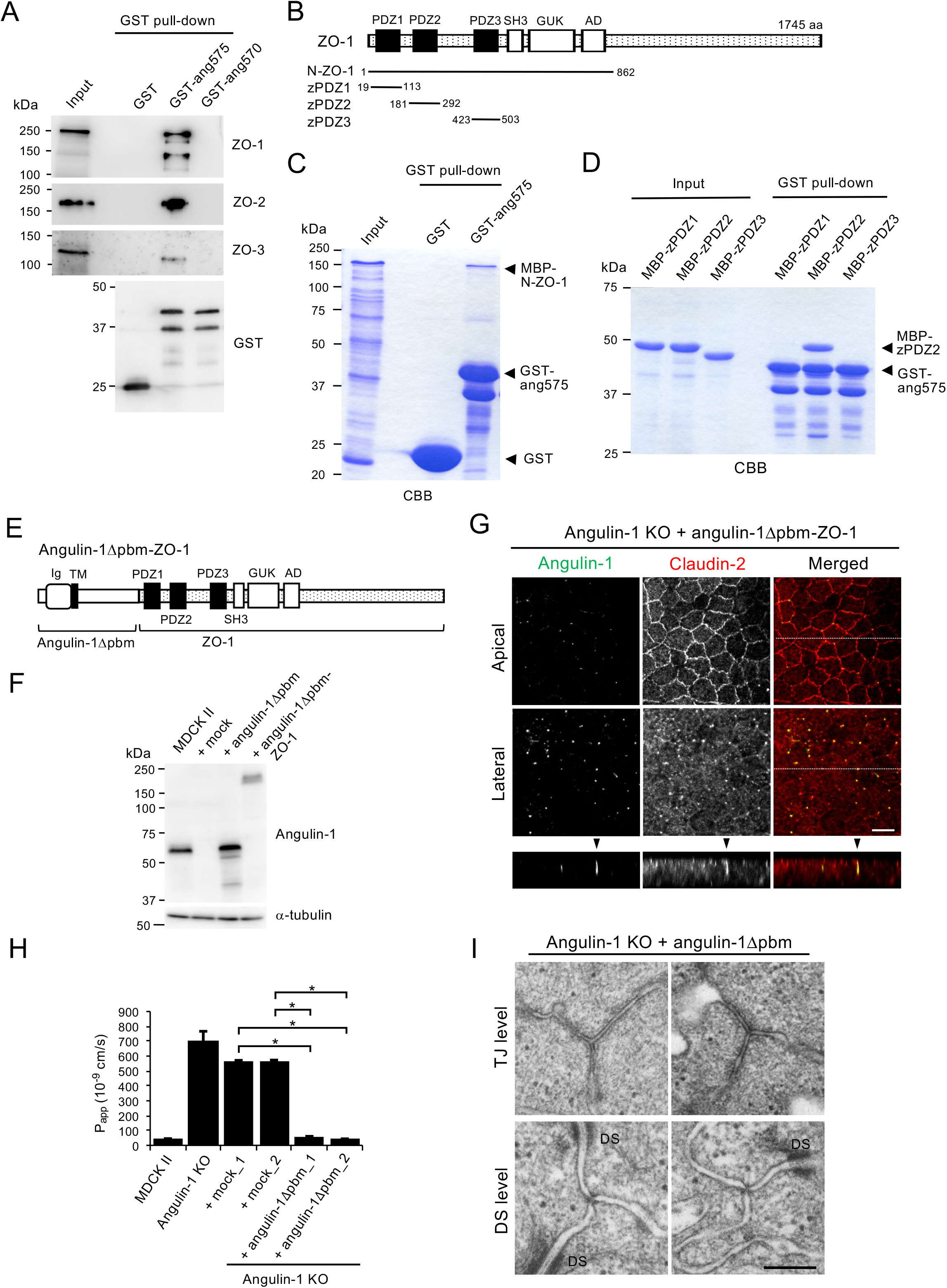
Direct binding of the angulin-1 pbm to ZO-1 and its role in tTJ formation. **(A)** Lysates of angulin-1 KO cells were incubated with GST and GST-fusion proteins of aa 409-575 and aa 409-570 of angulin-1 (GST-ang575 and GST-ang570, respectively), and subjected to GST pull-down assays. The lysates (input) and precipitates with GST, GST-ang575, and GST-ang570 were analyzed by western blotting with anti-ZO-1 mAb, anti-ZO-2 pAb, and anti-ZO-3 pAb. The precipitates were also immunoblotted with anti-GST pAb. **(B)** Schematic diagram of the domain structure of mouse ZO-1 protein with 1745 amino acids. ZO-1 contains three PDZ domains, an SH3 domain, a guanylate kinase domain (GUK), and an acidic domain (AD) in its N-terminal half. Four distinct portions of ZO-1 indicated as solid lines with amino acid numbers were produced as recombinant fusion proteins with MBP. **(C)** GST pull-down assays of MBP-N-ZO-1 fusion protein with GST-ang575. Input: crude lysate of *Escherichia coli* expressing MBP-N-ZO-1. **(D)** GST pull-down assays of MBP-fusion proteins of PDZ1, PDZ2, and PDZ3 of ZO-1 (MBP-zPDZ1, MBP-zPDZ2, and MBP-zPDZ3, respectively) with GST-ang575. Input: each purified MBP-fusion protein. In (C) and (D), the samples were subjected to SDS-PAGE followed by Coomassie Brilliant Blue (CBB) staining. **(E)** Schematic diagram of a chimeric protein of angulin-1 Δpbm and ZO-1 (angulin-1 Δpbm-ZO-1). **(F)** Western blotting of lysates of MDCK II cells, mock-transfected angulin-1 KO cells, and angulin-1 KO cells expressing angulin-1 Δpbm or angulin-1 Δpbm-ZO-1 with anti-angulin-1 pAb or anti-α-tubulin mAb. **(G)** Angulin-1 KO cells expressing angulin-1 Δpbm-ZO-1 were immunostained with anti-angulin-1 mAb and anti-claudin-2 mAb. Confocal sections of the apical region including claudin-2 staining and the lateral region together with the corresponding Z-stack images along the white dotted lines are shown. Arrowheads indicate TCs. **(H)** Paracellular flux of fluorescein in MDCK II cells, angulin-1 KO cells, two independent mock-transfected angulin-1 KO cell clones (+mock_1 and +mock_2), and two independent angulin-1 KO cell clones expressing exogenous angulin-1 Δpbm (+angulin-1Δpbm_1 and +angulin-1Δpbm_2). Data are shown as mean ± SD (*n*=3) and were analyzed by the Tukey–Kramer test. \**P*<0.01. The data in MDCK II cells, angulin-1 KO cells, and two independent mock-transfected angulin-1 KO cell clones are identical to those in Fig. 3 E. **(I)** EM observation of horizontal ultrathin sections of angulin-1 KO cells expressing exogenous angulin-1 Δpbm. TCs at the level of TJs (TJ level) and the more basal level of the lateral membrane containing desmosomes (DS level) were analyzed. DS: desmosome. Bars: 10 μm (G) and 200 nm (I).

We further examined the roles of angulin-1 pbm in the epithelial barrier function and plasma membrane contact formation at TCs. As shown in Figure 6 H, angulin-1 Δpbm-expressing angulin-1 KO cells had significantly decreased flux of fluorescein than control angulin-1 KO cells. Consistently, introduction of angulin-1 Δpbm to angulin-1/claudin-2 double KO cells increased TER and suppressed paracellular flux of fluorescein (Fig. S2, E–G). In horizontal ultrathin sections of angulin-1 Δpbm-expressing angulin-1 KO cells, the extracellular space in TCs at not only the TJ level but also the desmosome level was sealed by the plasma membranes (Fig. 6 I). These results suggest that the angulin-1 pbm is not essential for the epithelial barrier function or plasma membrane contact at TCs.

### The plasma membrane contact at TCs occurs independently of claudins and JAM-A

Angulin-1 Δpbm showed extended localization along the apicobasal axis unaccompanied by claudin-2 when expressed in angulin-1 KO cells (Fig. 5 B), but was able to seal the extracellular space at TCs. This suggests that angulin-1 may form the plasma membrane contact at TCs without claudins. To examine this possibility, we analyzed the morphology of TCs in claudin-based TJ strand-deficient epithelial cells, which we recently established by genome editing-based disruption of claudin-1, −2, −3, −4, and −7 genes in MDCK II cells (claudin quinKO cells) (Otani et al., 2019). Claudin quinKO cells lack TJ strands, but retain JAM-A mediated plasma membrane appositions with ~6-7 nm distance at the apical junctional complex level (Otani et al., 2019). Immunofluorescence staining revealed that angulin-1 was localized at the apical region, but not the lateral region, of TCs in claudin quinKO cells (Fig. 7 A). EM observation of horizontal ultrathin sections revealed that the extracellular space between the three plasma membranes at TCs was obliterated in claudin quinKO cells at the apical cell-cell junction level with plasma membrane appositions (Fig. 7 B). The plasma membrane contact at TCs was also observed in horizontal sections containing desmosomes (Fig. 7 B).

**Figure 7.**
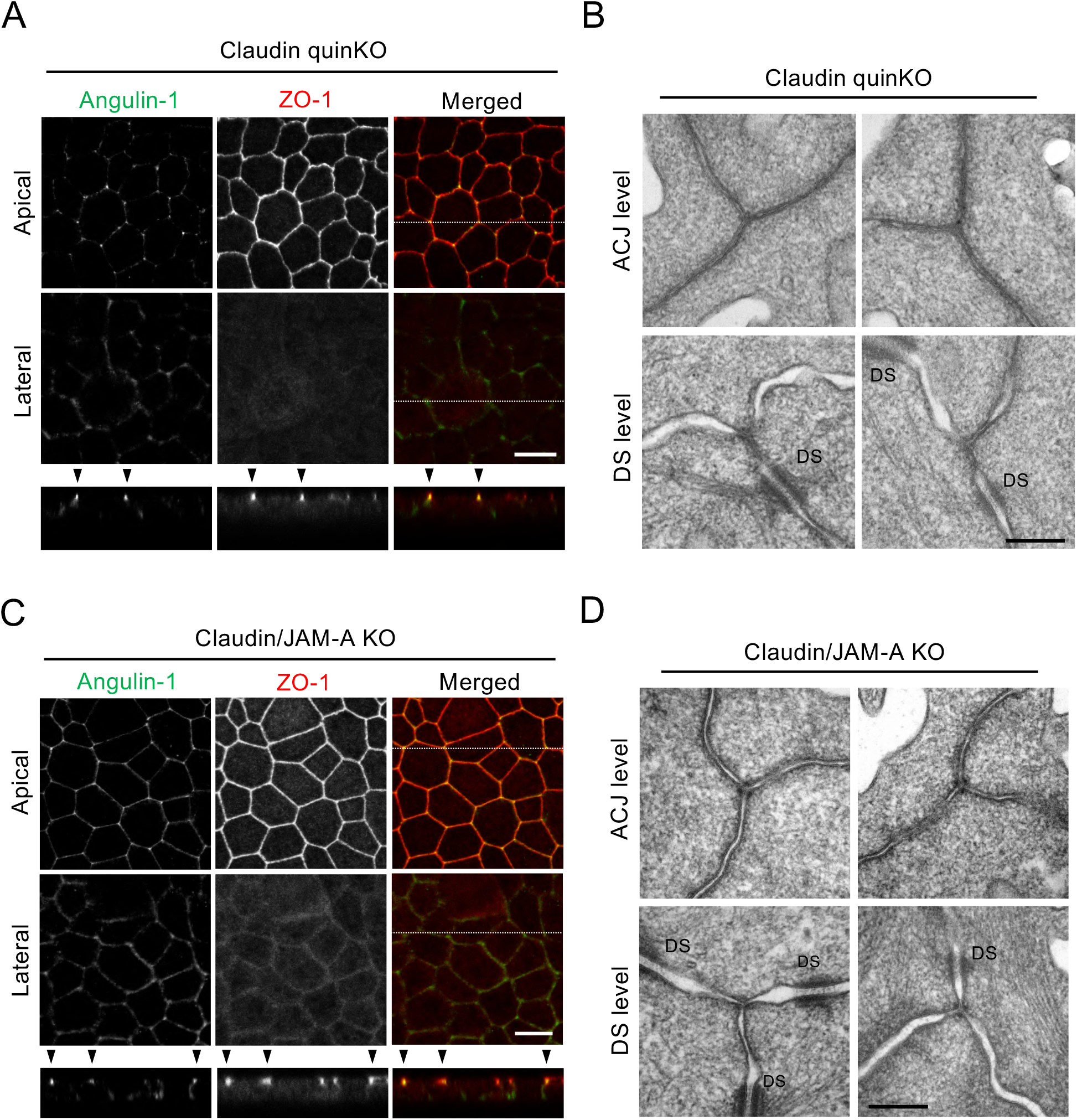
The plasma membrane contact at TCs is maintained in claudin quinKO cells and claudin/JAM-A KO cells. **(A)** Immunofluorescence staining of claudin quinKO cells with anti-angulin-1 pAb and anti-ZO-1 mAb. **(B)** EM observation of horizontal ultrathin sections of claudin quinKO cells. **(C)** Immunofluorescence staining of claudin/JAM-A KO cells with anti-angulin-1 pAb and anti-ZO-1 mAb. **(D)** EM observation of horizontal ultrathin sections of claudin/JAM-A KO cells. In (A) and (C), confocal sections of the apical region including ZO-1 staining and the lateral region together with the corresponding Z-stack images along the white dotted lines are shown. In both cells, signals for angulin-1 localized at TCs were detected in the apical region only. Arrowheads indicate TCs. Bars: 10 μm. In (B) and (D), TCs at the level of apical cell-cell junctions without desmosomes (ACJ level) and the more basal level of the lateral membrane containing desmosomes (DS level) were analyzed. Note that the intercellular space is widened in the apical region of claudin/JAM-A KO cells. DS: desmosome. Bars: 200 nm.

To examine the requirement of JAM-A-mediated membrane appositions for angulin-1 assembly and plasma membrane contact formation at TCs, we analyzed claudin/JAM-A KO cells established from claudin quinKO cells by disrupting the JAM-A gene (Otani et al., 2019). Angulin-1 was localized at the only apical part of TCs on immunofluorescence staining of claudin/JAM-A KO cells (Fig. 7C). On EM of horizontal ultrathin sections of the cells, the plasma membrane contact at TCs was clearly observed in sections containing parallel plasma membranes with a narrow space, which appeared to the apical cell-cell junction level. The plasma membrane contact was also observed in sections containing desmosomes (Fig. 7D). These results demonstrate that neither claudin-based TJ strands nor JAM-A-mediated membrane appositions are required for the plasma membrane contact at TCs.

### Tricellulin is required for connection of TJ strands to the central sealing elements, but not for epithelial barrier function

Considering that angulins recruit tricellulin to TCs, the angulin-1-mediated plasma membrane contact at TCs may be attributed to tricellulin. To examine this idea, we established four independent tricellulin-deficient cell clones from MDCK II cells using CRISPR/Cas9-mediated genome editing (Figs 8, A, B and S3). Tricellulin contains four transmembrane domains, N- and C-terminal cytoplasmic domains, two extracellular loops, and a short cytoplasmic turn. The obtained tricellulin KO cell clones may allow expression of the N-terminal half of tricellulin with a frameshift around aa 275, located at the cytoplasmic turn between the second and third transmembrane domains. However, no signals for truncated tricellulin at TCs were detected in these cells by immunofluorescence staining with an antibody against the N-terminal cytoplasmic region of tricellulin (Fig 8 B), indicating that tricellulin was functionally impaired. Immunofluorescence staining of tricellulin KO cells revealed that angulin-1 was localized at TCs with apicobasal extension and colocalized with claudin-2 (Fig. 8 C). ZO-1 was mostly colocalized with angulin-1 along the apicobasal axis at TCs, but was infrequently missing from some angulin rods (Fig. 8 C). Unlike angulin-1 KO cells (Fig. 3 D), all four tricellulin KO cell clones showed no increase in paracellular flux of fluorescein compared with MDCK II cells (Fig. 8 D). Moreover, alterations in TER and paracellular flux of fluorescein were not detected in tricellulin KO cells on a claudin-2 KO background (Fig. S4, A–D). Finally, we analyzed the TC ultrastructure in tricellulin KO cells by EM. Horizontal ultrathin sections at the TJ level revealed that tricellulin KO cells contained either the plasma membrane contact or a gap at TCs (Fig. 8 E). The plasma membrane contact at TCs was also observed at the desmosome level in tricellulin KO cells (Fig. 8 E). On freeze-fracture replica EM, the central sealing elements with apicobasal extension were observed at TCs in tricellulin KO cells. Intriguingly, however, there were hardly any connections of short TJ strands to the central sealing elements (Fig. 8, F_1_, F_1_’, F_2_, and F_3_). At some TCs, apical TJ strands from both sides appeared to cave in and attach to one another to form central sealing elements that extended in the basal direction (Fig. 8 F_3_). The cave-in of TJ strands likely corresponded to the gap at TCs at the TJ level observed in horizontal ultrathin sections. These observations suggest that tricellulin is required for the organization of tTJs by connecting short TJ strands to the central sealing elements, but is not essential for the formation of the central sealing elements and plasma membrane contact at TCs or the epithelial barrier function.

**Figure 8.**
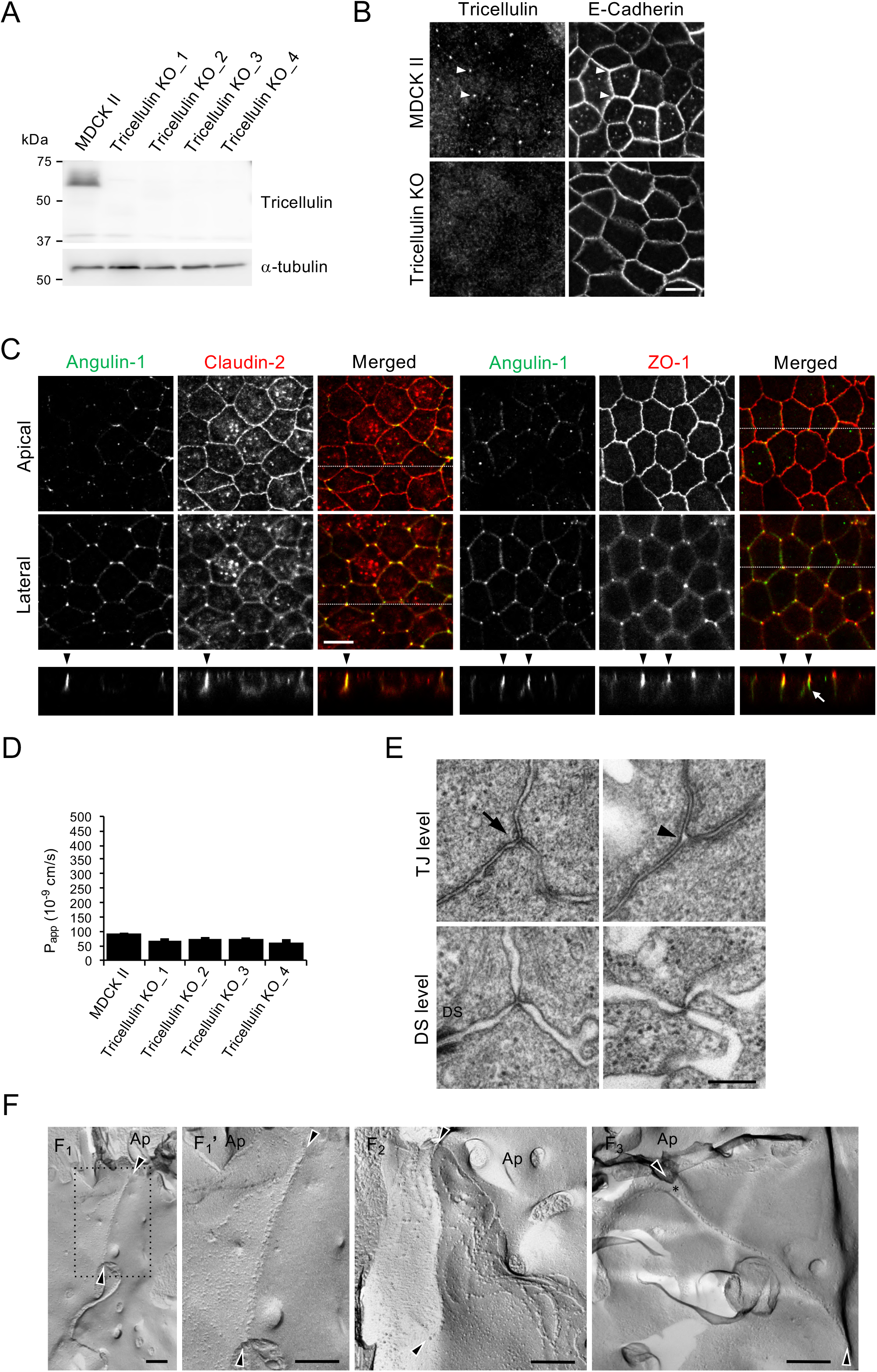
Roles of tricellulin in tTJ formation and epithelial barrier function. **(A)** Lysates of four independent tricellulin KO cell clones (tricellulin KO_1–4) were analyzed by western blotting with anti-tricellulin pAb or anti-α-tubulin mAb. **(B)** Tricellulin KO_4 cells were immunostained with anti-tricellulin pAb and anti-E-cadherin mAb. Arrowheads indicate TCs. **(C)** Double immunofluorescence staining of tricellulin KO_4 cells with anti-angulin-1 pAb and anti-claudin-2 mAb or anti-angulin-1 pAb and anti-ZO-1 mAb. Arrowheads indicate TCs. Confocal sections of the apical region including TJ markers and the lateral region together with the corresponding Z-stack images along the white dotted lines are shown. The white arrow indicates an angulin-1-positive region at a TC without ZO-1. **(D)** Paracellular flux of fluorescein in MDCK II and tricellulin KO_1-4 cells. Data are shown as mean ± SD (*n*=3). **(E)** EM observation of horizontal ultrathin sections of tricellulin KO_4 cells. TCs at the level of TJs (TJ level) and the more basal level of the lateral membrane containing desmosomes (DS level) were analyzed. The TJ level in tricellulin KO cells contained either the plasma membrane contact (arrow) or a gap (arrowhead) at TCs. Close plasma membrane contacts at TCs were also observed at the DS level. **(F)** Freeze-fracture EM observation of tricellulin KO_4 cells. Arrowheads indicate the central sealing elements without connection of short TJ strands. The asterisk indicates a cave-in of vertical TJ strands. Ap: apical side. Bars: 10 μm (B and C) and 200 nm (E and F).

## Discussion

In the present study, we examined the roles of angulin-1 and tricellulin in tTJ formation and epithelial barrier function in MDCK II cells by loss-of-function analyses using genome editing. We found that angulin-1 is required for the plasma membrane contact at TCs, central sealing element formation, and epithelial barrier function, while tricellulin is not (Fig. 9). We also found that claudin-based TJ strands are dispensable for the plasma membrane contact at TCs. These results suggest that angulin family proteins are essential for the plasma membrane seal at TCs, which should be a key step in tTJ formation, independently of tricellulin and claudins.

**Figure 9.**
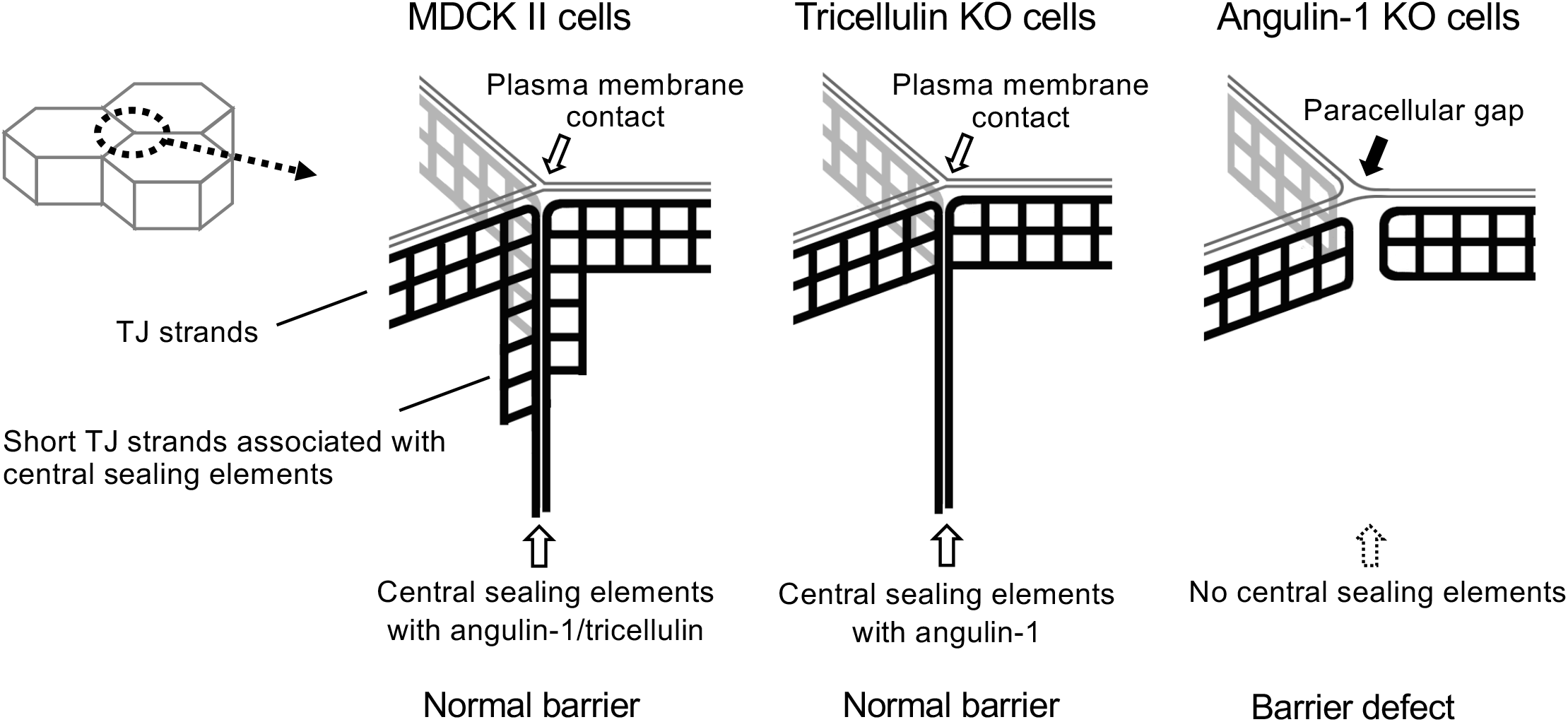
Summary of the phenotypes of MDCK II, angulin-1 KO, and tricellulin KO cells for tTJ formation and epithelial barrier function. In MDCK II cells, angulin-1 and tricellulin were localized at TCs extending along the apicobasal axis. The plasma membrane contact at TCs was observed in horizontal ultrathin sections. In freeze-fracture replicas, typical tTJs containing the central sealing elements and associated short TJ strands were observed. In tricellulin KO cells, angulin-1 was localized at TCs extending along the apicobasal axis. The plasma membrane contact and the central sealing elements were maintained, but the central sealing elements lacked connection of short TJ strands. In angulin-1 KO cells, a paracellular gap was observed at TCs in horizontal ultrathin sections. In freeze-fracture replicas, the vertical TJ strands at TCs were separated and did not form the central sealing elements. The epithelial barrier function was impaired in angulin-1 KO cells, but not in tricellulin KO cells.

### Apicobasal extension of tTJs

We found that tTJ proteins, including tricellulin and angulin family proteins, extended along the apicobasal axis at TCs in mouse nephrons and MDCK II cells by immunofluorescence microscopy. Because tricellulin and angulin-1 are localized along the central sealing elements of tTJs on freeze-fracture immunolabeling EM (Ikenouchi et al., 2005)(Masuda et al., 2011), it is reasonable that the rod-like fluorescent signals of tTJ proteins reflect the outline of tTJs by light microscopy. In mouse proximal tubules, tricellulin signals were visualized as dots, rather than rods, suggesting that the tTJs in proximal tubule epithelial cells along the apicobasal axis are relatively short. In rats, it was reported that TJs at bicellular contacts in proximal tubules contain 1–2 TJ strands, while those in collecting ducts contain ~7 TJ strands (Claude and Goodenough, 1973)(Schiller et al., 1980). The thickness of TJs may influence the depth of tTJs along the apicobasal axis. Importantly, claudins, occludin, and ZO-1 were distributed along the apicobasal axis at TCs and colocalized with tricellulin and angulins, suggesting that TJ-associated proteins are also incorporated into tTJs. Ikenouchi et al. (2005) reported that tricellulin was detected as vertically oriented rods at TCs, while occludin was concentrated as dots at the most apical region of the tricellulin rods in Z-stack confocal sections of mouse EpH4 mammary epithelial cells evaluated by immunofluorescence. These descriptions cannot exclude the existence of occludin in tTJs, because the TJ-to-tTJ ratio of occludin may vary depending on cell types. Considering the limitations of light microscopy, it is difficult to clarify whether these TJ-associated proteins are localized at the central sealing elements and/or short TJ strands connected to the central sealing elements within tTJs by immunofluorescence staining. In tricellulin KO cells, the central sealing elements were observed in tTJs by freeze-fracture EM, and claudin-2 was colocalized with angulin-1 along the apicobasal axis at TCs by immunofluorescence staining. These observations suggest that claudins are incorporated into the central sealing elements of tTJs. Further investigations by super-resolution microscopy or freeze-fracture immunolabeling EM are needed to reveal the precise distributions of TJ-associated proteins within tTJs.

### Roles of ZO-1 binding to angulin-1 in tTJ formation

We found that the pbm in the angulin-1 C-terminus directly binds to PDZ2 of ZO-1. The pbm-deleted angulin-1 mutant was distributed along the apicobasal axis at TCs and colocalized with tricellulin, but not ZO-1 and claudins. However, forced conjunction of ZO-1 to the pbm-deleted angulin-1 mutant in angulin-1 KO cells induced apicobasal extension of claudins at TCs. These results suggest that ZO-1 mediates claudin recruitment to angulin-1 along the apicobasal axis for tTJ formation. Consistent with this idea, PDZ1 of ZO-1 can directly bind to the C-terminal pbm of claudins (Itoh et al., 1999). However, the real molecular mechanism for TJ protein assembly in tTJs appears more complex. In tricellulin KO cells, claudin-2 was colocalized with angulin-1 along the apicobasal axis at TCs. Consistently, the central sealing elements were observed in these cells by freeze-fracture EM. However, ZO-1 was incompletely colocalized with angulin-1 along the apicobasal axis at TCs in tricellulin KO cells. These observations indicate that tricellulin is involved in the colocalization of ZO-1 with angulin-1 and that ZO-1 is not the only determinant for colocalization of claudins with angulin-1. Riazuddin et al. (2006) observed direct binding of the N-terminal half of ZO-1 to the C-terminal cytoplasmic domain of tricellulin by *in vitro* binding assays. Meanwhile, Ikenouchi et al. (2008) showed that a claudin-3 mutant lacking the C-terminal pbm and tricellulin were highly colocalized at cell-cell contacts when overexpressed in mouse L fibroblasts without recruiting ZO-1, suggesting an interaction between claudins and tricellulin in the plasma membrane regardless of ZO-1. It was also shown that claudins lacking the C-terminal pbm can polymerize in the plasma membrane to form TJ strand-like structures in mouse L fibroblasts (Furuse et al., 1999)(Ikenouchi et al., 2008), suggesting an intrinsic ability of claudins for TJ strand formation at least under certain conditions. These molecular interactions may cooperatively regulate the recruitment of claudins to angulin-1-positive regions at TCs to form tTJs.

### Roles of angulins in the plasma membrane contact at TCs

As the most important finding in this study, we demonstrated that angulin-1 is involved in the plasma membrane contact at TCs to obliterate the paracellular space. The plasma membrane contact at TCs was disrupted in angulin-1 KO cells, but maintained in tricellulin KO cells. Moreover, the plasma membrane contact at TCs was observed in not only claudin quinKO cells lacking claudin-based TJ strands, but also claudin/JAM-A KO cells lacking JAM-A-mediated plasma membrane appositions. The pbm-deleted angulin-1 mutant lacking ZO-1-binding ability generated the plasma membrane contacts at TCs when expressed in angulin-1 KO cells. Taken together, angulin-1 is responsible for the plasma membrane contact at TCs independently of tricellulin, claudins, JAM-A, and binding to ZO-1. In angulin-1 KO cells, the reduction in epithelial barrier function corresponded to gap formation at the paracellular space of TCs, while continuous TJ strands at bicellular contacts were observed in freeze-fracture replicas. These observations indicate that angulin-1-mediated obliteration of the paracellular space at TCs is crucial for the full barrier function of epithelial cellular sheets. It is necessary to establish whether angulin-2 and angulin-3 can generate the plasma membrane contact at TCs and the central sealing elements at tTJs. If these proteins can reconstitute tTJs in angulin-1 KO cells similar to angulin-1, it will be of interest to clarify the functional differences among the angulin subtypes.

In claudin quinKO cells and claudin/JAM-A KO cells, angulin-1 was localized at apical TCs, which merged with the apical cell-cell junction marker ZO-1, but was not present at lateral TCs, which did not merge with ZO-1. However, the plasma membrane contact at TCs was observed in horizontal ultrathin sections of the cell layers, even at the desmosome level corresponding to the lateral TCs. In claudin quinKO cells and claudin/JAM-A KO cells, angulin-1 may extend basally and form the plasma membrane contact to some extent, although it may not be detectable by confocal microscopy with poor Z-axis resolution. Alternatively, a trace amount of angulin-1 may extend to the lateral TCs and form the plasma membrane contact there. The depth of the plasma membrane contact at TCs in these cells requires further clarification by serial ultrathin section analyses.

Even in the absence of claudins and JAM-A, the extracellular space at the plasma membrane contact at TCs appeared to have almost no gap, similar to TJs in ultrathin section EM. When expressed in mouse L fibroblasts lacking cell adhesion activity, angulin-1 sporadically assembled in cell-cell contacts between adjoining cells, suggesting that angulin-1 has weak homophilic interactions (Masuda et al., 2011). However, it remains unclear whether angulin-1 itself has strong adhesive activity to cross-link the plasma membranes at TCs. However, considering that JAM-A with two Ig-like domains in its extracellular region generates very close membrane appositions with only 6–7-nm distance (Otani et al., 2019), it may be reasonable that angulin-1 with a single Ig-like domain can form the plasma membrane contact at TCs with almost no extracellular space.

The present observations indicate that the basic mechanisms for bicellular TJ formation and tTJ formation are independent of one another, but are coordinately organized in epithelial cells to form a functional paracellular barrier. Recently, giant unilamellar vesicles containing reconstituted claudin-4 were shown to form adhesive membrane interfaces with a fence function against extracellular membrane proteins (Belardi et al., 2018). In these vesicles, close membrane contacts were absent at trivesicular junctions, indicating that claudins are not sufficient for tTJ formation. It will interesting to examine whether angulins can cross-link trivesicular junctions using this simple reconstitution system in which the lipid content can be controlled, to understand the mechanism for angulin-mediated membrane contact formation at TCs.

### Roles of tricellulin in tTJ formation

In our study, the plasma membrane contact at TCs and epithelial barrier function were retained in tricellulin KO cells. Thus, the question arises as to the role of tricellulin in tTJs. In freeze-fracture EM, tricellulin KO cells had basally extended central sealing elements at TCs, but lost short TJ strands connected to the central sealing elements. Similar structural defects in tTJs were observed in the inner ear epithelia of Tric^R497X/R497X^ mice, which mimic a mutation in the human tricellulin-encoding gene causing hearing loss (Nayak et al., 2013). These observations indicate that tricellulin is required for connection of short TJ strands to the central sealing elements of tTJs. Consistent with this notion, tricellulin expression increased the number of end-to-side connections in TJ strands reconstituted by claudin-1 in mouse L fibroblasts (Ikenouchi et al., 2008) and HEK293 cells (Cording et al., 2013), resulting in the formation of compressed TJ strand meshworks. However, the mechanism by which tricellulin induces end-to-side connections of TJ strands remains elusive. We previously showed that the N-terminal cytoplasmic domain of tricellulin binds to the Cdc42 GEF Tuba, activates Cdc42, and regulates F-actin organization during cell-cell junction formation in cultured epithelial cells (Oda et al., 2014). It will be of interest to elucidate how tricellulin regulates juxtamembrane F-actin organization at TCs through Cdc42 activation and whether this action of tricellulin influences end-to-side connections of TJ strands. Alternatively, tricellulin may mediate end-to-side connections of TJ strands as a joint. Recently, Van Itallie et al. (2017) showed that occludin tends to concentrate at TJ strand ends and end-to-side junction points of TJ strands in Rat-1 fibroblasts expressing claudins and occludin (Van Itallie et al., 2017). Considering that tricellulin shares structural similarity with occludin (Ikenouchi et al., 2005), tricellulin may concentrate at the junctions of TJ strands and facilitate their branching.

Despite the loss of short TJ strands in the vicinity of the central sealing elements, we did not detect any impairment of the epithelial barrier function in tricellulin KO cells by conventional barrier assays. This suggests that tricellulin has a subtle role in the epithelial barrier function. It was reported that RNAi-mediated suppression of tricellulin decreased TER and increased paracellular flux in mouse EpH4 mammary epithelial cells (Ikenouchi et al., 2005), while overexpression of tricellulin reduced paracellular flux in MDCK II cells (Krug et al., 2009). The discrepancy between these previous studies and the present study may arise from differences in the cells or experimental conditions used. Meanwhile, the notion of a subtle role for tricellulin in the epithelial barrier function appears consistent with the results of *in vivo* loss-of-function studies on tricellulin. No significant decreases in endocochlear potential were observed in Tric^R497X/R497X^ mice and tricellulin KO mice, although the mice showed progressive hearing loss associated with hair cell degeneration (Nayak et al., 2013)(Kamitani et al., 2015). In Tric^R497X/R497X^ mice and tricellulin KO mice, a biotin-based tracer did not permeate from the perilymph to the stria vascularis (Nayak et al., 2013)(Kamitani et al., 2015). Furthermore, although tricellulin is ubiquitously expressed in epithelial tissues, no other clinical manifestations were co-segregated with hearing loss in two DFNB49 families with tricellulin mutations (Nayak et al., 2015). In contrast, the phenotypes caused by angulin deficiencies are diverse and severe. Angulin-1-deficient mice exhibit embryonic lethality with blood-brain barrier failure (Mesli et al., 2004)(Sohet et al., 2015), while angulin-2-deficient mice show polyuria and polydipsia arising from renal concentrating defects in addition to hearing loss (Higashi et al., 2015)(Morozko et al., 2015)(Sang et al., 2015)(Gong et al., 2017). This evidence supports the idea that angulins, but not tricellulin, play essential roles in the plasma membrane seal at TCs and epithelial barrier function.

## Materials and Methods

### Cell culture and antibodies

MDCK II cells were provided by Dr. Masayuki Murata (The University of Tokyo, Tokyo, Japan). Cells were grown in DMEM (low glucose; #05919; Nissui) supplemented with 10% fetal bovine serum. Rat anti-tricellulin mAb (C96) (Ikenouchi et al., 2005), rat anti-occludin mAb (Saitou et al., 1997), rabbit anti-angulin-1 pAb (Oda et al., 2020), mouse anti-ZO-1 mAb (Itoh et al., 1991), rabbit anti-occludin pAb (Saitou et al., 1997), rat anti-angulin-1 mAb (Iwamoto et al., 2013) and rabbit anti-tricellulin pAb (Oda et al., 2014) were generated as described previously. Mouse anti-claudin-2 mAb (32-5600), rabbit anti-tricellulin pAb (48-8400), rabbit anti-ZO-2 pAb (38-9100) and rabbit anti-ZO-3 pAb (36-4100) were purchased from Thermo Fisher Scientific. Rat anti-E-cadherin mAb (ECCD2; M108, Takara Bio), mouse anti-α-tubulin mAb (14-4502-82, eBioscience) and goat anti-GST pAb conjugated with HRP (RPN1236, Amersham Biosciences) were obtained commercially. Alexa 488-conjugated donkey anti-rabbit, mouse or rat IgG was obtained from Thermo Fisher Scientific. Cy3-conjugated donkey anti-rabbit, mouse or rat IgG was purchased from Jackson ImmunoResearch Laboratories. Horseradish peroxidase (HRP)-linked anti-rabbit, mouse or rat IgG was purchased from GE Healthcare.

### Expression vectors and transfection

The cDNA encoding mouse angulin-1 of 575 amino acids was described previously (Higashi et al., 2013). To construct expression vectors for full-length angulin-1 and angulin-1 Δpbm, cDNAs encoding mouse angulin-1 and its mutant lackning the C-terminal 5 amino acids were amplified by PCR using KOD-Plus-Ver.2 DNA polymerase (Toyobo) and subcloned into pCAGGSneodelEcoRI (Niwa et al., 1991). To construct expression vectors for GST-fusion of aa 409-575 and aa 409-570 of mouse angulin-1, the corresponding cDNAs of mouse angulin-1 were amplified by PCR using KOD-Plus-Ver.2 DNA polymerase and subcloned into pGEX-6P-1 (GE Healthcare). To construct an expression vector for MBP-tagged N-ZO-1 (aa 1-862), cDNA encoding aa 1-862 of mouse ZO-1 was amplified by PCR using KOD-Plus-Ver.2 DNA polymerase and subcloned into pMAL-cRI (New England BioLabs). Expression vectors for MBP-fusion proteins of PDZ1, PDZ2, and PDZ3 domains of ZO-1 (Itoh et al., 2001) were kindly provided by Dr. Masahiko Itoh (Dokkyo Medical University, Tochigi, Japan). To construct an expression vector for a fusion protein of angulin-1 Δpbm with ZO-1, the BglII sites within the angulin-1 Δpbm cDNA and ZO-1 cDNA were disrupted in advance without changing amino acid sequences by inverse PCR. The cDNA encoding angulin-1 Δpbm was amplified by PCR using KOD-Plus-Ver.2 DNA polymerase and subcloned into pCAGGSneodelEcoRI between NotI site and BglII site. Next, cDNA encoding ZO-1, in which BglII site was disrupted, was amplified by PCR using KOD-Plus-Ver.2 DNA polymerase and the DNA fragments were subcloned into pCAGGSneodelEcoRI between BglII site and EcoRI site.

DNA transfection was performed using the Lipofectamine LTX and Plus Reagent (Thermo Fisher Scientific) according to the manufacturer’s instructions. Stable cell lines were selected by treatment with 500 μg/ml G-418 (Nacalai Tesque).

### Genome editing

Angulin-1 KO cells were established from parental MDCK II cells by genome editing using TALEN. TALENs were constructed according to the instruction provided by the TALE Toolbox kit from the Zhang laboratory (Sanjana et al., 2012) (Addgene, #1000000019). The target sequences for the left arm, spacer and right arm are indicated in Fig S1. The construction of the TALEN expression vectors and transfection were performed as described previously (Tokuda et al., 2014). Transfected cell colonies were propagated and screened for angulin-1 depletion by immufluorescence staining. Angulin-1 KO cell clones were isolated by limiting dilution of cell colonies containing angulin-1-negative cells. To confirm mutations in the targeting sites in angulin-1 gene in angulin-1 KO cells, the genome DNA containing the targeting site was amplified by PCR using a primer set, 5’-GCCCTTTAACGTCCTGGGAC-3’ (forward) and 5’-GAGCAACTCCTCTCACTCCG-3’ (reverse), subcloned into pTAC-1 vector (BioDynamics Laboratory Inc.) by TA-cloning, and subjected to Sanger sequencing.

Tricellulin KO cells were generated from parental MDCK II cells with Cas9-gRNA RNP complexes using a CUY21 Pro-Vitro electroporator (Nepagene). CRISPR RNA for the target sequence, 5’-CGGCATGACCACCTACTACC<u>GGG</u>-3’ (PAM site is underlined), and trans-activating CRISPR RNA were synthesized by IDT and annealed with each other. Cas9 proteins (IDT) were incubated with the gRNA duplex at room temperature for 10 min to form Cas9-gRNA RNP complexes (1:1.2 molar ratio). 100 pmol Cas9 and 120 pmol gRNA duplex were introduced into 1×10^6^ cells. Electroporation was performed with the following conditions: 150 V for 10 msec (prepulse), 10 pulses of 20 V for 50 msec at 50 msec intervals (postpulses). Angulin-1/claudin-2 double KO cells and tricellulin/claudin-2 double KO cells were established from claudin-2 KO cells (Tokuda and Furuse, 2015) by CRISPR/Cas9-mediated genome editing of angulin-1 gene and tricellulin gene, respectively, using pSpCas9(BB)-2A-Puro(PX459) V2.0 vector (62988; Addgene). The following DNA sense and antisense strands of the targeting sites were annealed with each other in KOD FX neo buffer (Toyobo): angulin-1, 5’-CACCGGCTGGGGCGCGGTCGTCTT-3’ (sense) and 5’-AAACAAGACGACCGCGCCCCAGCC-3’ (antisense); tricellulin, 5’-CACCGCGGCATGACCACCTACTACC-3’ (sense) and 5’-AAACGGTAGTAGGTGGTCATGCCGC-3’ (antisense). The obtained DNA duplexes were ligated to pSpCas9(BB)-2A-Puro(PX459) V2.0 digested by BpiI (FD1014; Thermo Fisher Scientific). Claudin-2 KO cells were transfected with the CRISPR/Cas9 vectors using Lipofectamine LTX and Plus Reagent (Thermo Fisher Scientific). The transfected cells were cloned in glass bottom 96 well plate (Corning) by limiting dilution and the KO cells were selected by immunofluorescence microscopy. To confirm mutations in the corresponding targeting sites, the genomic regions of targeting sequences were amplified by PCR with primer sets of SalI site-containing forward primers and EcoRI site-containing reverse primers. The PCR products were digested with SalI/EcoRI and subcloned into SalI/EcoRI-digested pBluescript SK (-). Mutations in targeting sites were confirmed by Sanger sequencing. The following primers were used for genomic PCR: angulin-1, 5’-GGGGTCGACCGGAGCGGAGGCGGGAAGGGGAGG-3’ (forward) and 5’-GGGGAATTCCGGCGGTGGGGACTCCATCCATCG-3’ (reverse); tricellulin, 5’-GGGGTCGACGAGCAGCGAGCGGGAGGAGGACTTGC-3’ (forward) and 5’-GGG GAATTCCCACCTCGTGCCTCCACAGCTTCAGG-3’ (reverse).

### Immunofluorescence microscopy

For immunofluorescence microscopy of cultured cells, cells were seeded at a density of 1.0×10^5^ cells/cm^2^ on 12 mm-diameter transwell filters with 0.4 μm pore size (3401; Corning). After 3-4 days of culture on transwell filters, cells were fixed with 1% formaldehyde in PBS containing 0.5 mM CaCl_2_ for 10 min at room temperature and washed with PBS three times. The cells were permeabilized with 0.2% Triton X-100 in PBS for 10 min at room temperature and washed with PBS three times. The samples were blocked with 1% BSA in PBS for 30 min at room temperature. The cells were incubated with primary antibodies followed by fluorescence-labeled secondary antibodies. After washing with PBS, the samples were embedded in FluoroSave reagent (Millipore) and observed with a laser scanning confocal microscope (TCS-SPE; Leica Microsystems) equipped with a HCX PL APO 63×/1.40 objective. Images were processed using Fiji/ImageJ software (National Institutes of Health).

### Electron microscopy

For ultrathin section electron microscopy of cultured cells, samples were prepared in principle as described previously (Otani et al., 2019). Cells were seeded at a density of 1.0×10^5^ cells/cm^2^ on 12 mm-diameter Transwell polycarbonate filters (3401; Corning). After 3-4 days of culture, the cells were fixed with a fixative containing 2% glutaraldehyde, 2% paraformaldehyde, 20 mM CaCl_2_ and 0.1 M cacodylate buffer (pH7.4) for 1 h at room temperature and washed with 0.1 M cacodylate buffer (pH7.4). Filters were excised by scalpels and post-fixed with 1% OsO_4_ in 0.1 M cacodylate buffer, pH7.4, for 1 h on ice. After washing with water, the samples were stained with 0.5% uranyl acetate for 2 h at room temperature. After washing with water, the samples were dehydrated in a graded series of ethanol. The samples were then soaked in propylene oxide for 1-2 min, transferred to a 1:1 mixture of propylene oxide and Quetol 812 resin and incubated overnight. The samples were embedded in Epon 812 resin. Ultra-thin sections of ~60 nm thickness were stained with 0.5% uranyl acetate for 3 min and then stained with Sato’s lead solution containing 1% lead citrate, 1% lead nitrate and 2% sodium citrate (Sato, 1968) for 3 min. The sections were washed with water and dried. For freeze-fracture electron microscopy, cells were seeded at a density of 1.0×10^5^ cells/cm^2^ on 24 mm-diameter Transwell polycarbonate filters (3412; Corning). After 3-4 days of culture, the cells were fixed with 2% glutaraldehyde in 0.1 M phosphate buffer (pH7.4) for 1 h and washed three times with 0.1 M phosphate buffer (pH7.4). The samples were immersed in 30% glycerol in 0.1 M phosphate buffer (pH7.4) for 30 min at room temperature and then frozen with liquid nitrogen. The frozen samples were fractured at −100°C and platinum-shadowed unidirectionally at an angle of 45° using a Balzers freeze etching system (BAF060; Bal-Tec). The fractured samples were collected on formvar-filmed grids. Samples were observed with a transmission electron microscope (JEM-1011 or JEM-1010; JEOL) at an accelerating voltage of 80 kV.

### Western blotting and GST pull-down assay

For Western blotting of cell lysates, cells were lysed with Laemmli SDS sample buffer supplemented with 100 mM DTT and boiled at 100°C for 5 min. The proteins were separated by SDS-PAGE using a 10, 12.5 or 15% polyacrylamide gel and transferred to PVDF membrane with 0.45 μm pore size (Millipore). The membranes were blocked with 5% skim milk in TBS supplemented with 0.1% Tween 20 (TBS-T) and incubated with primary antibodies diluted with 5% skim milk in TBS-T or immunoreaction enhancer solution of Can Get Signal (NKB-101; Toyobo) for 1 h at 37°C. After the incubation, the membranes were washed three times with TBS-T, followed by incubation of HRP-linked secondary antibodies diluted with 5% skim milk in TBS-T or immunoreaction enhancer solution of Can Get Signal (NKB-101; Toyobo) for 1 h at 37°C. After the incubation, the membranes were rinsed three times with TBS-T. The secondary antibodies were detected by enhanced chemiluminescence (ECL Prime; GE Healthcare). Images were obtained with LAS3000 mini (Fujifilm) and processed using Fiji/ImageJ software (National Institutes of Health).

For *in vitro* binding assays, all of GST-tagged or MBP-tagged proteins were expressed in *Escherichia coli* (DH5α). For GST pull-down assays using lysates of angulin-1 KO cells, angulin-1 KO cells were lysed with the lysis buffer (0.5% NP-40, 50 mM Tris-HCl, pH7.4, 150 mM NaCl, 2 mM EDTA, 10% glycerol) supplemented with 1 mM DTT and protease inhibitor cocktail (Nacalai Tesque). After centrifugation of the lysates, the supernatant fluids were incubated with Glutathione Sepharose 4B beads coupled with GST or GST-angulin-1 cytoplasmic region (aa 409-575) for 2-3 h at 4°C. The beads were washed three times with the lysis buffer and boiled with 2×Laemmli SDS sample buffer supplemented with 100 mM DTT. For in vitro binding assays between angulin-1 and ZO-1, MBP-tagged proteins of ZO-1 (N-ZO-1: 1-862 aa, PDZ1 domain: 19-113 aa, PDZ2 domain: 181-292 aa or PDZ3 domain: 423-503 aa) were incubated with GST or GST-tagged angulin-1 cytoplasmic region (409-575 aa or 409-570 aa) for 2-3 h at 4°C and further incubated with Glutathione Sepharose 4B beads (GE Healthcare) for 1 h at 4°C. The beads were washed three times with the lysis buffer and boiled with 2×Laemmli SDS sample buffer supplemented with 100 mM DTT. The lysates were analyzed by SDS-PAGE and Western blotting.

### TER and paracellular tracer flux

MDCK II cells were seeded at a density of 1.0×10^5^ cells/cm^2^ on 12 mm-diameter transwell polycarbonate filters with 0.4 μm pore size (3401; Corning) and cultured for 4-5 days. Electrical resistance was measured using Millicell ERS-2 (Millipore). Electrical resistance of transwell filters without cells was measured as a blank. The mean blank value was subtracted from electrical resistance and TER was determined by multiplying the electrical resistance by the growth area of the transwell filter. To measure paracellular tracer flux, medium was changed to phenol-red free DMEM (08489-45; Nacalai Tesque) supplemented with 10% fetal bovine serum and 2 mM L-glutamine (16948-04; Nacalai Tesque). On the following day, 200 μM fluorescein (16106-82; Nacalai Tesque) was added to the apical chamber. Medium volume of the apical and the basal chamber was 250 μl and 1 ml, respectively. The cells were incubated for 2 h at 37°C and 5% CO_2_. The medium of the basal chamber was collected and the fluorescent intensity of the medium was measured by a microplate reader (SpectraMax Paradigm; Molecular Devices). Fluorescent intensity of medium without fluorescent tracer was measured as a blank. After subtraction of the mean blank values from fluorescent intensities of the samples, the apparent permeability (Papp) was calculated using the following equation: P_app_=(dQ/dt)/AC_o_ (dQ: the amount of tracer transported to the basal chamber; dt: incubation time; A: the area of the transwell filters; C_o_: initial concentration of tracer in the apical chamber).

## Acknowledgments

We thank Shinsaku Tokuda for designing TALEN constructs, Masayuki Murata and Masahiko Itoh for kindly providing reagents, the EM facility in the National Institute for Physiological Sciences for support in EM, Osamu Nagata, Mika Watanabe, and Yuichiro Kano for technical assistance, Motohiro Nishida for use of a microplate reader, and Akira Nagafuchi, Shigenobu Yonemura, and all members of the Furuse laboratory for discussions. We also thank Alison Sherwin, PhD, from Edanz Group (https://en-author-services.edanzgroup.com/) for editing a draft of this manuscript. This work was supported by a JSPS Grant-in-Aid for Challenging Exploratory Research (16K15226), JSPS Grants-in-Aid for Scientific Research (B) (26291043 and 18H02440), and a Life Science Aid from the Takeda Science Foundation to MF.

