## Supplementary Figures for "Angulin-1 seals tricellular contacts independently of tricellulin and claudins"

A

R G W G A V V (aa 19-25)

GGTGGGCCCCGGCCGGCCTCTCGGGGCTGGGGCGCGGTCGTCTTCGGGTGCCTCTTTCTCAGC

Left arm — Spacer — Right arm

KO_1 — Deletion: CGGGGCT–GGGCGCGGTCGT

KO_2 — Insertion: CGGGGCTGGGCGCGCGGTCGT

KO_2 — Insertion: CGGGGCTGGGGGCGCGGTCGT

B

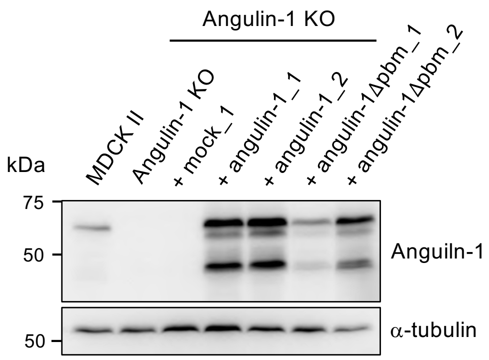


C

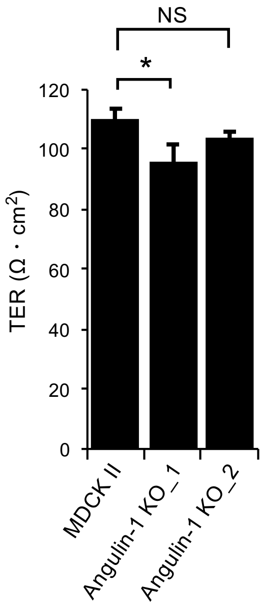


Supplementary Figure S1

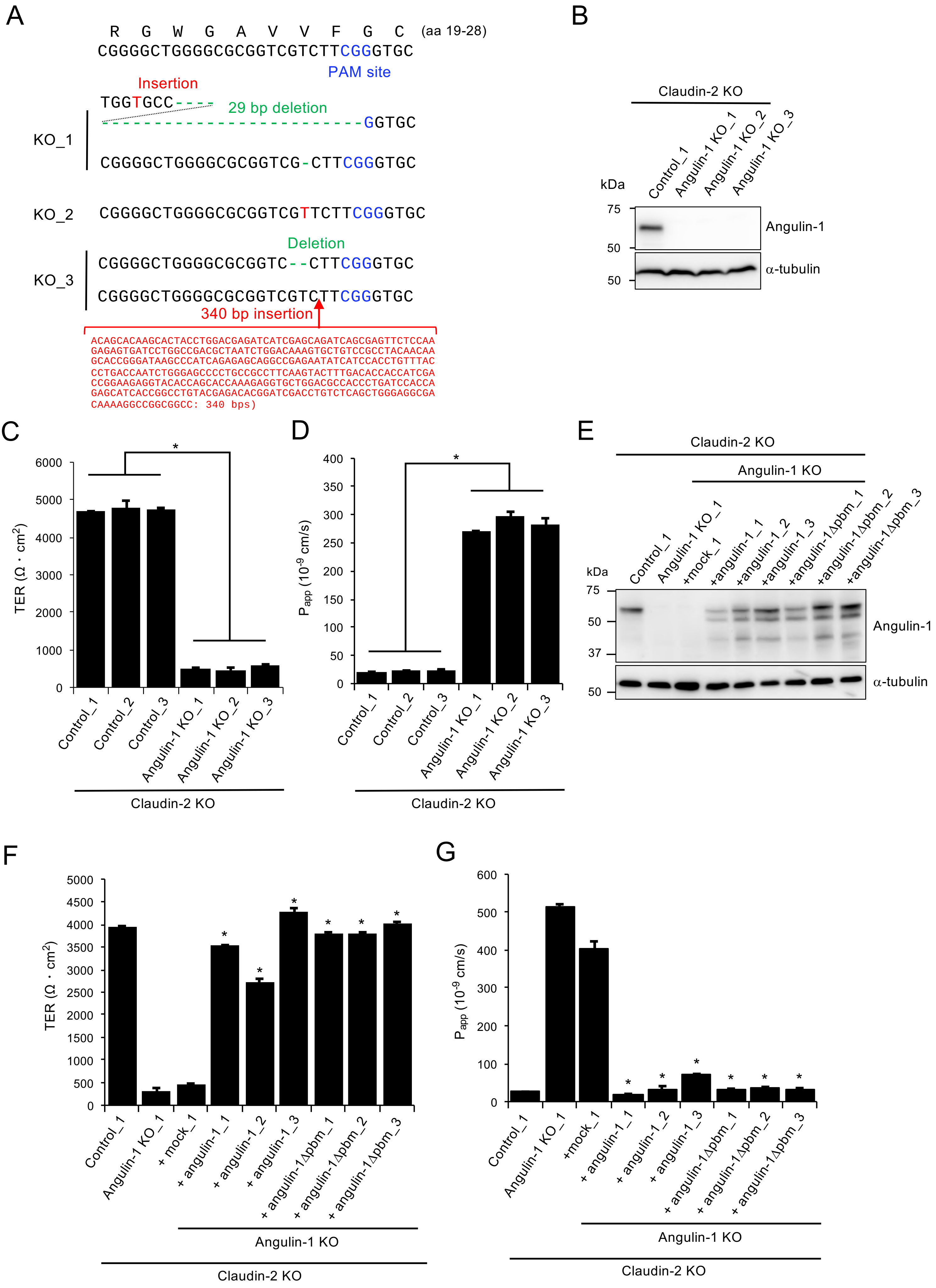


Supplementary Figure S2

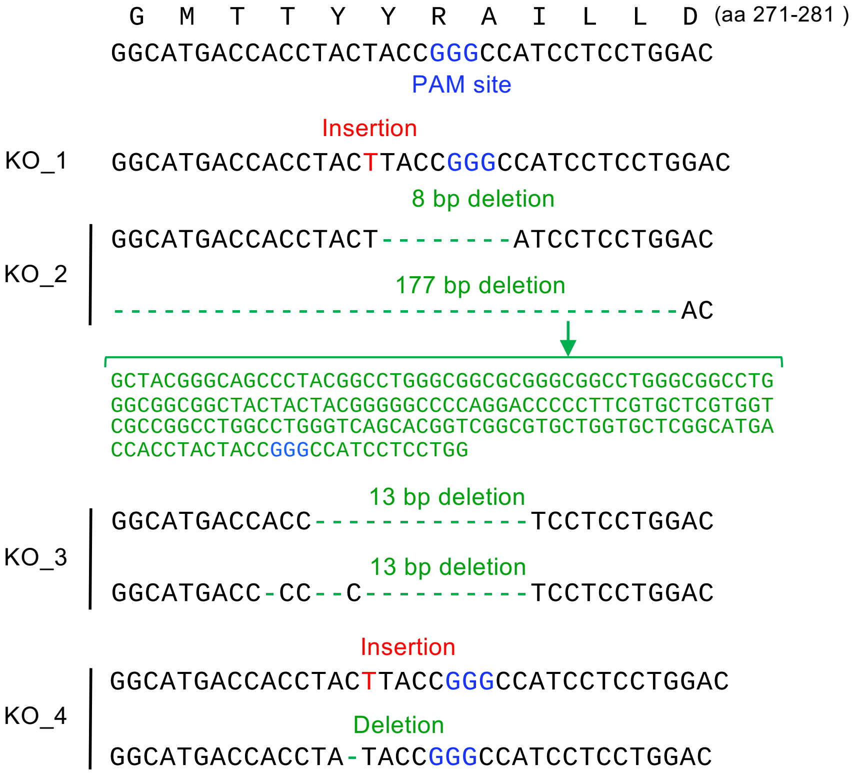


Supplementary Figure S3

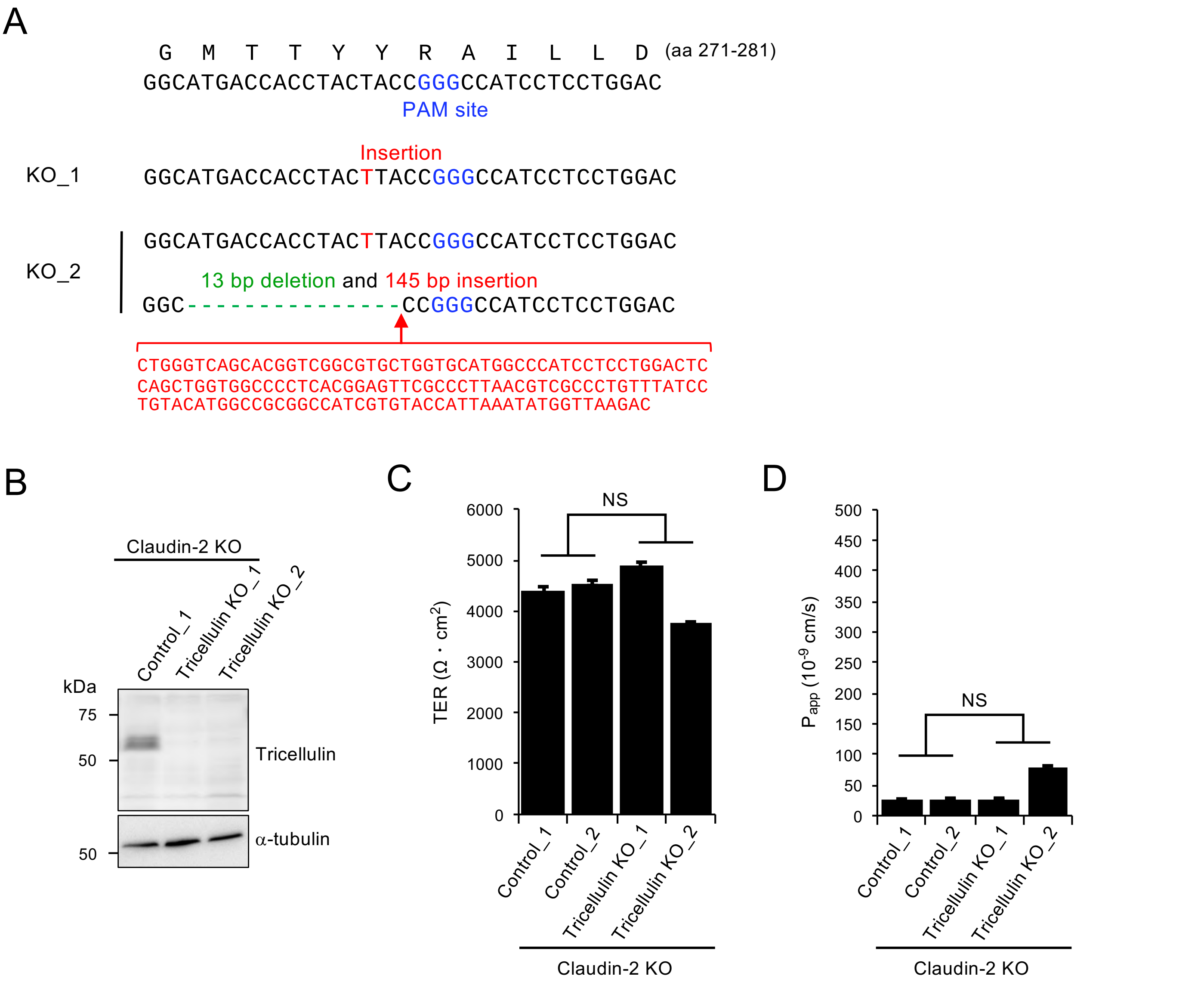


Supplementary Figure S4

**Supplementary Figure legends**

**Figure S1. Characterization of angulin-1 KO cells and their derivatives. (A)** Establishment of angulin-1 KO cells by TALEN-mediated genome editing. The left and right arms of the TALEN targeting sites indicated in blue were set at the sides of the 20-nt spacer sequence encoding aa 19–25 of dog angulin-1. The DNA sequence of dog angulin-1 was obtained from the Ensemble genome database (ENSCAFT00000011359.4). DNA sequencing revealed that frameshift mutations were introduced in the spacer sequence in two angulin-1 KO cell clones (KO_1 and KO_2). **(B)** Western blotting of lysates of MDCK II cells, angulin-1 KO cells, a mock-transfected angulin-1 KO cell clone, two angulin-1 KO cell clones expressing exogenous angulin-1, and two angulin-1 KO cell clones expressing angulin-1Δpbm with anti-angulin-1 pAb or anti-α-tubulin mAb. **(C)** TER measurements of MDCK II cells and two angulin-1 KO cell clones. Data were analyzed by Dunnett's test ($n$=3 for MDCK II cells vs. $n$=3 for each angulin-1 KO cell clone). Data are shown as mean ± SD ($n$=3). *$P$<0.05.

**Figure S2. Characterization of angulin-1/claudin-2 double KO cells and their derivatives. (A)** Frameshift mutations introduced upstream of the PAM site in the angulin-1 gene in angulin-1/claudin-2 double KO cell clones generated by CRISPR/Cas9-mediated genome editing. **(B)** Western blotting of lysates of a control claudin-2 KO cell clone and three angulin-1/claudin-2 double KO cell clones with anti-angulin-1 pAb or anti-α-tubulin mAb. **(C and D)** TER (C) and paracellular flux of fluorescein (D) were measured in three control claudin-2 KO cell clones and three angulin-1/claudin-2 double KO cell clones. Data for TER and paracellular flux of fluorescein were analyzed by Student's $t$-test and Welch's $t$-test, respectively ($n$=9 total for three control clones [$n$=3 per clone] vs. $n$=9 total for three angulin-1/claudin-2 double KO clones [$n$=3 per clone]). **(E)** Western blotting of lysates of a control claudin-2 KO cell clone, an angulin-1/claudin-2 double KO cell clone, a mock-transfected angulin-1/claudin-2 double KO cell clone, three angulin-1/claudin-2 double KO cell clones expressing exogenous angulin-1, and three angulin-1/claudin-2 double KO cell clones expressing exogenous angulin-1Δpbm with anti-angulin-1 pAb or anti-α-tubulin mAb. **(F and G)** TER (F) and paracellular flux of fluorescein (G) were measured in the stable cell clones shown in (E). Data were analyzed by Dunnett's test ($n$=3 for the mock cell clone vs. $n$=3 for each angulin-1- or angulin-1Δpbm-expressing clone). Data in (C), (D), (F), and (G) are shown as mean ± SD ($n$=3). *$P$<0.01.

**Figure S3. Characterization of tricellulin KO cells**

Frameshift mutations were introduced in the proximity of the PAM site in the tricellulin gene in four tricellulin KO clones (KO_1–4) established by CRISPR/Cas9-mediated genome editing.

**Figure S4. Characterization of tricellulin/claudin-2 double KO cells**

**(A)** Frameshift mutations introduced in the proximity of the PAM site in the tricellulin gene in two tricellulin/claudin-2 double KO cell clones (KO_1 and KO_2) established by CRISPR/Cas9-mediated genome editing. **(B)** Western blotting of lysates of a control claudin-2 KO cell clone and two tricellulin/claudin-2 double KO cell clones with anti-tricellulin pAb or anti-α-tubulin mAb. **(C and D)** TER (C) and paracellular flux of fluorescein (D) were measured in two control claudin-2 KO cell clones (Control_1 and 2) and two tricellulin/claudin-2 double KO cell clones (KO_1 and KO_2). Data are shown as mean ± SD ($n$=3) and were analyzed by Welch's $t$-test ($n$=6 total for two control clones [$n$=3 per clone] vs. $n$=6 total for two tricellulin/claudin-2 double KO clones [$n$=3 per clone]).
